# Modeling Attention and Binding in the Brain through Bidirectional Recurrent Gating

**DOI:** 10.1101/2024.09.09.612033

**Authors:** Saeed Salehi, Jordan Lei, Ari S. Benjamin, Klaus-Robert Müller, Konrad P. Kording

## Abstract

Attention is a key component of the visual system, essential for perception, learning, and memory. Attention can also be seen as a solution to the binding problem: concurrent attention to all parts of an entity allows separating it from the rest. However, the rich models of attention in computational neuroscience are generally not scaled to real-world problems and there are thus many behavioral and neural phenomena that current models cannot explain. Here, we propose a bidirectional recurrent model of attention that is inspired by the emerging understanding of biological object-based attention and modern neural networks for image segmentation. It conceptualizes recurrent connections as a multi-stage internal gating process where bottom-up connections transmit features, while top-down and lateral connections transmit attentional gating signals. Our model can recognize and segment simple stimuli such as digits as well as objects in natural images and is able to be prompted with object labels, attributes or locations. It can learn to perform a range of behavioral findings, such as object binding, selective attention, inhibition of return, and visual search. It also replicates a variety of neural findings, including increased activity for attended objects, features, and locations, attention-invariant tuning, and relatively late onset attention. Most importantly, our proposed model unifies decades of cognitive and neurophysiological findings of visual attention into a single principled architecture. Our results highlight that the ability to selectively and dynamically focus on specific parts of stimulus streams can help artificial neural networks to better generalize and align with human brains.

## Introduction

*Attention* is a mechanism by which the brain selects a meaningful subset of incoming stimuli for perception, learning, or memory [Carrasco, 2011; Hommel et al., 2019; Itti et al., 2005; Lindsay, 2020; Maunsell, 2015; Moore and Zirnsak, 2017; Tsotsos, 2021]. By focusing on a subset of the world, attention promises a more localized, sparse, and less noisy signal in response to sensory input [Posner et al., 1980]. For learning, focusing on the relevant subset of stimuli (i.e., a target) leads to better generalization [Rutishauser et al., 2004; Walther et al., 2005]. In the case of memory, sparser and cleaner signals could be reused more often in the stimulus set (e.g., objects or object parts) and thus be more efficiently stored, recalled, and compositionally utilized [Kruijne et al., 2021; Olivers and Roelfsema, 2020]. Hence, attention plays an indispensable role in neural information processing and understanding attention is critical to advancing our comprehension of learning, memory and perception.

What makes attention difficult to model is the widespread nature of the various phenomena it includes. Attention appears on different scales and through different mechanisms [Buschman and Kastner, 2015; Itti et al., 2005; Maunsell, 2015; Moore and Zirnsak, 2017; Posner, 2016]. One frequently studied form is spotlight attention, where focus is directed towards a region in the environment, enhancing perception and processing of information in that region [Itti and Koch, 2001; Wolfe and Horowitz, 2004]. Behavioral experiments show that perception in such attention foci is faster and more reliable [Chun and Wolfe, 2005; Posner, 1980]. Another frequently studied form is feature-based attention where, as with space, knowledge about features of relevant stimuli speeds up perception and lowers error rates [Bichot et al., 2015; Treisman, 1998; Treisman and Gelade, 1980]. A third form of attention, arguably related to feature-based attention, is object-based attention, where attention is bound to an object as a whole, including its spatial and visual features [Baldauf and Desimone, 2014; Behrmann et al., 1998; Scholl, 2001; Treisman, 1998]. A fundamental question here is how an object that is represented by activity across millions of neurons and many cortical regions can be perceived and attended to as a single object [Von der Malsburg, 1999]. This is known as the binding problem, asking how the brain can integrate and merge all of the activities relating to the same entity and how to separate them from all the other entities [Feldman, 2013; Robertson, 2003; Singer, 2001; Treisman, 1996; Von der Malsburg, 1995]. Hence, understanding object-based attention is a crucial step in solving the feature binding problem [Roelfsema, 2023; Treisman, 1998]. Thus, neuroscientists and psychologists have long studied the relationship between attention and binding and sought to build a unifying model of these phenomena.

Despite their theoretical elegance, typical models of biological attention lack one or more elements that are crucial for building a comprehensive model of object-based attention [Cavanagh et al., 2023; George et al., 2020; Posner, 2023]. The first element is the top-down process, where activity in higher-level brain areas modulates the neural activity in lower-level cortices and has been shown to influence whole-object perception and integration [Freeman et al., 2003; Gilbert and Li, 2013; Paneri and Gregoriou, 2017; Thiele and Bellgrove, 2018]. The second element is lateral processes and communication, shown to be relevant for binding, noise reduction, perception of continuity and closure, and many more [Field et al., 1993; George et al., 2020; Gray et al., 1989; Han et al., 2005]. A third element is recurrence, where activity travels in all directions simultaneously and involves repeated processing of information over time [Hochreiter and Schmidhuber, 1997]. Indeed, there are strong experimental and theoretical evidence for the importance of recurrence in visual processing, attention, and binding [Kar et al., 2019; Kietzmann et al., 2019; O’Reilly et al., 2013; Roelfsema, 2023; Thorat et al., 2021; van Bergen and Kriegeskorte, 2020], and has led to considerable interest in recurrent models for attention [Kubilius et al., 2019; Larsen and Druckmann, 2022; Linsley et al., 2020; Mnih et al., 2014]. For all the richness of the existing literature, we are still lacking a model that integrates all of the aforementioned elements, is capable of handling natural stimuli, and at the same time describes the many known findings in the field.

Attention has also been studied and utilized in machine learning, albeit often under different interpretations of the concept. Among them, the transformer architecture stands out as the most popular deep learning architecture bearing the name of attention. Transformers [Vaswani et al., 2017] use self-attention to dynamically create a sparse and weighted combination of input feature representations. But despite their immense success and ability to learn powerful representations [Caron et al., 2021; Dosovitskiy et al., 2020; Kirillov et al., 2023], transformers lack the three key elements of attention: lateral, top-down, and recurrent activities, which restricts their biological plausibility and limits their ability to solve tasks that require understanding of multiscale features to produce precise local binding. Similarly, other deep learning models of attention either lack recurrency [Kaul et al., 2019], or top-down control and internal gating [J. Ba et al., 2014; Locatello et al., 2020; Mnih et al., 2014; Nayebi et al., 2018], or use decoupled mechanisms for feature learning and attention [Stollenga et al., 2014]. Nonetheless, deep learning has been remarkably successful in object recognition, segmentation, and representation learning [Bengio et al., 2013; George et al., 2017; LeCun et al., 2015; Ronneberger et al., 2015] and thus promises new approaches for neuroscientists to build effective and generalizable models of visual attention [Richards et al., 2019].

Here, we propose a deep learning model of biological attention, by means of *bottom-up, top-down*, and *lateral connections* operating through a *bidirectional recurrent gating* mechanism. Our model is inspired by the emerging understanding of object-based attention, while borrowing elements and techniques from state-of-the-art machine learning. We base our architecture on the U-Net [Ronneberger et al., 2015], while incorporating *residual connections* [He et al., 2016], *divisive normalization* [J. L. Ba et al., 2016; Reynolds and Heeger, 2009], attention-driven *neuromodulation*, and task prompts as explicit input. The network incorporates a novel multitask-embedding paradigm as a tool to further improve its flexibility and generalization. We show that our model can learn to perform a broad range of attention tasks, including but not limited to: object recognition, segmentation, spatial binding, inhibition of return, border ownership, and top-down visual search. Additionally, we illustrate the importance of recurrent attention in solving feature-binding problems using a pathological dataset and and its extension to real-world scenarios. We later scale up the network to handle attention tasks on complex natural stimuli, such as those from the COCO dataset. Finally, we demonstrate how our model describes known neurophysiological phenomena, including attention-invariant tuning, attention-mediated neuromodulation, and late-onset attention, effectively unifying a broad range of behavioral and neural phenomena related to object-based attention.

## Results

We aimed to develop a biologically motivated model that could simultaneously handle real-world naturalistic stimuli and solve benchmark tasks commonly used by cognitive scientists and neuroscientists to study vision and attention in animals and humans. Our architecture consists of a feedforward contracting *Feature* pathway that hierarchically processes and extracts learned features, alongside a top-down expansive *Attention* pathway that combines lateral and top-down connections to implement task- and stimulus-dependent bidirectional recurrent gating mechanism (Fig. 1 a, see supplementary materials). The lateral connections form a long-range bidirectional recurrent network, while the dense recurrent layers provide the model with working memory and thus, enabling it to perform memory-dependent tasks and behaviors such as inhibition of return. Since recurrency is central to our architecture, the model expects a sequence of images as its input stimulus. Correspondingly, it outputs a sequence of attention maps and label predictions, generated sequentially in each iteration. Besides the input stimuli, the network can also accept additional inputs, such as the task index for multitask learning and prompts for top-down tasks. Altogether, our model can effectively learn to perform multiple tasks on the same architecture (Fig. 1 b).

**Figure 1.**
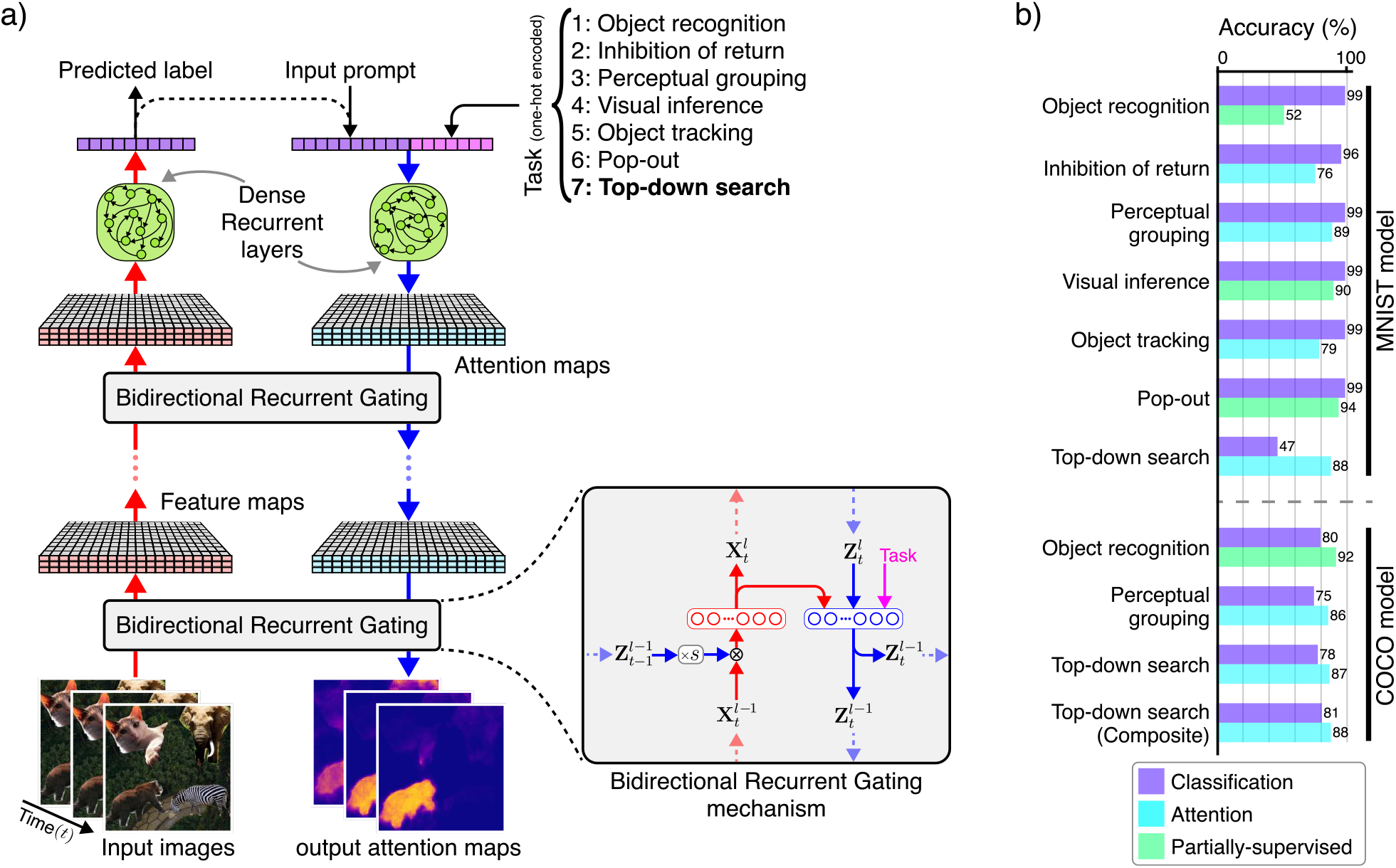
Model architecture and multitask learning. **a)** Bidirectional recurrent gating is the core block of our model. In this figure, we use color-coding to denote the bottom-up and top-down information streams and processes, with red representing bottom-up and blue representing top-down. Subscripts indicate the corresponding time (i.e., iteration) and superscripts denote the corresponding layer. The network comprises of two main pathways: the *Feature* pathway, which hierarchically extracts learned feature representations (shown in red); the *Attention* pathway, which combines top-down information, the task signal, and feature maps to generate attention maps (shown in blue). The attention maps consequently and multiplicatively modulate the feature maps of the next iteration. The two pathways meet at the bottleneck, which incorporates the dense recurrent and linear layers and outputs estimated label. In addition to the bottleneck, the two pathways communicate feature maps and attention maps through the lateral connections. **b)** Our architecture enables effective multitask learning on both simple (i.e., digits from MNIST) and complex (i.e., animals from COCO dataset) stimuli. Here we show the classification and attention accuracy for the two models trained on COCO and MNIST compositions (see see supplementary materials for detailed architectures). For some tasks, we use partially-supervised training, meaning that we only provide one supervision signal during training, either target attention maps (i.e., segmentation) or target labels (i.e., classification).

We use stochastic gradient descent [Rumelhart et al., 1986] to train the model on a combination of two objectives: (a) the network should be able to classify objects, for which we use Cross-Entropy loss, and (b) the network should be able to segment objects in the input layer, for which we use Mean-Squared-Error loss. Although at the first glance these two goals seem insufficient for learning complex tasks, we will demonstrate that our model can leverages the multitask paradigm to effectively learn partially-supervised tasks and attend to the correct visual features. For regularization, we use *l*^2^-norm for learnable weights. During inference, we measure the accuracy, both for classification and attention maps, and pixel error for the final iteration (see supplementary materials).

### Behavioral results and multitask learning

Visual attention encompasses an extensive range of behavioral findings [Itti et al., 2005], including: a) object recognition in a cluttered scene, b) spatial cueing of attention and perceptual grouping, c) feature saliency and pop-out, d) top-down control, e) inhibition of return, and f) pre-attentive features, to name a few. In what follows, we design a series of multitask experiments to examine whether our network is able to replicate these known behavioral findings. The first experiment (arguably using toy data) is performed on compositions of handwritten digits from the MNIST dataset [LeCun, 1998] with various aspects extended to enable a broad set of tests. The second experiment is conducted on a rather complex set of stimuli based on the COCO dataset [Lin et al., 2014], though on fewer tasks (see the supplementary materials for single-task training results).

Object recognition is one of the primary objectives of the visual system, which requires separating the target object from the background and foreground and classifying it correctly. Although today most artificial models of object recognition rely simply on feedforward processes, the evidence from the brain implies a significant role played by attention and recurrency [DiCarlo et al., 2012; Super et al., 2010]. Attention and recurrent processing could reduce ambiguity of the edges shared between the target and other objects and improve accuracy and robustness [Fang et al., 2009; Konkle and Alvarez, 2024]. Moreover, the visual system tends to maintain a stable internal representation of the object even when the foreground or background moves. To test whether the model can solve these problems, we designed an object recognition and permanence task, where a digit is placed on top of a background and partially obscured by a foreground (Fig. 2 a). Both the background and foreground are randomly generated correlated noise that move slowly across the image, while the object remains stationary. Training for this task includes only the target label, requiring the network to learn to uncover a sensible attention map solely through classification error and multitask paradigm. Our model achieves 99% classification accuracy and pixel error of 0.02 in recovering the correct attention map in the final iteration on the test dataset. This demonstrates that the model can effectively solve object recognition, segmentation and permanence tasks in the presence of significant levels of correlated noise and occlusion.

**Figure 2.**
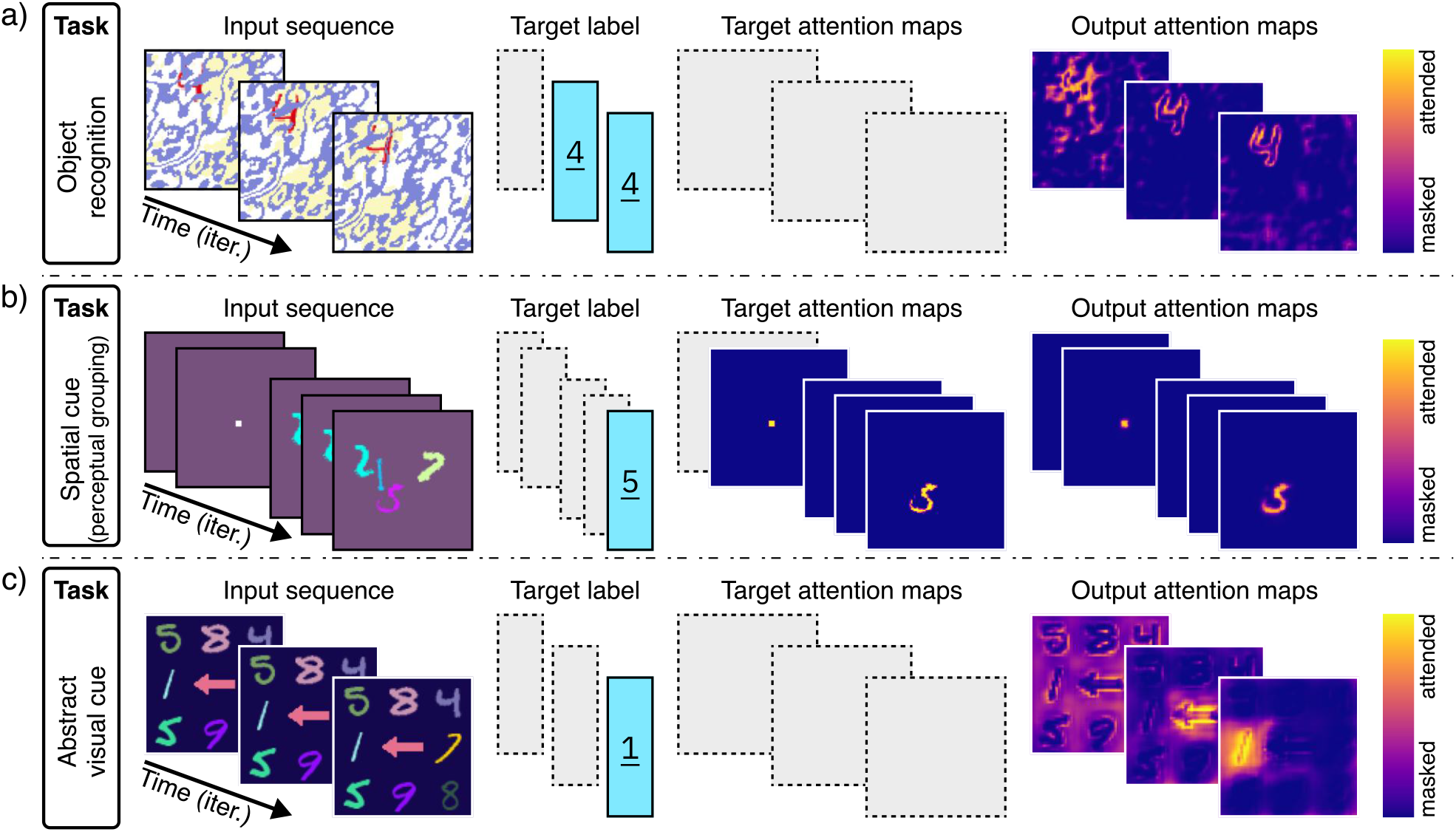
Multitask training on MNIST composites (Part 1/3). Results for a single model trained on seven tasks simultaneously. Here we present the results for: **a)** *object recognition*, **b)** *spatial cueing*, and **c)** *visual inference*. For each task, from left to right: the input sequence, target labels, target attention masks, and output (estimated) attention maps are shown. For target label and target attention map, the gray dashed boxes imply that the target is NOT provided during training. Colors for the task (a) input sequence have been inverted to improve visualization.

Spotlight attention is a well studied model system for attention [Chun and Wolfe, 2005; Itti and Koch, 2001]. A common element in cognitive visual experiments is to use a cue to spatially orient the attention to a region and task the subject to fixate on the cue and follow the instructions. Such cues could be also used to guide perceptual grouping [Herzog, 2018; Roelfsema, 2006]. Therefore, we tested our model to see if it could also learn where to direct its attention using spatial cues (Fig. 2 b). This was crucial as we intend to use a similar approach in the curve-tracing task later. Here, we trained the network to first attend to a cue (i.e., dot) appearing in first two iterations. The sequence is then followed by a random arrangement of few digits, and the network must attend to the digit that is contiguous with the previously shown cue. Our model achieves 99% classification accuracy and pixel error of 0.001 in recovering the correct attention map in the final iteration on the test dataset. These results demonstrate that our model possesses spotlight attention capabilities comparable to human subjects in such tasks.

Apart from spatial cues, we also use abstract visual cues to direct our attention to an object in the scene. The most prominent example would be pointing with the index finger. We have learned to first look at the finger, recognize the direction in which it is pointing towards, and then to orient our gaze and attention to the object in line with the finger’s direction. Therefore, we trained our model on such task, where 8 digits are arranged around an arrow and the goal is to locate and classify the target digit. During training, only information about the target digit, to which the arrow is pointing at, is provided (Fig. 2 c). It is important to note that since NO attention map for the arrow is given, the model has to learn about the arrow only through classification error and multitask paradigm. Our model achieves 99% in classification accuracy and pixel error of 0.05 in recovering the correct attention map in the final iteration on the test dataset. Interestingly, the attention maps indicate that the model has learned to attend to the arrow first to infer the location of the target digit, indicating that the model is using its knowledge acquired from other tasks that digits are separate entities from arrows, and therefore avoids learning the combination of all the arrow directions and digits, resulting in a more robust behavior. Thus, the system can learn about and attend to an abstractly cued location and use multitask paradigm for better generalization (see supplementary materials for our approach to circumvent this challenge in single-task learning).

Salient features in a scene are thought to be the main driving factors in pre-attention [Itti et al., 1998; Tsotsos et al., 1995; Wolfe, 2005; Zhaoping and Dayan, 2006], although the neural basis of this phenomenon is still debated [Einhäuser and König, 2003; Nothdurft, 2006; Parkhurst et al., 2002]. We have implemented a task where the model is presented with a grid of 9 digits, where all but one are of the same instance and color ((Fig. 3 a)). The model is trained to return the class of the digit that is different from the rest, without knowing the target attention map. The network successfully learns to detect and classify the salient digit with 99% classification accuracy and pixel error of 0.07 in recovering the correct attention map, thus indicating that our network can learn and perform saliency-dependent pop-out tasks.

**Figure 3.**
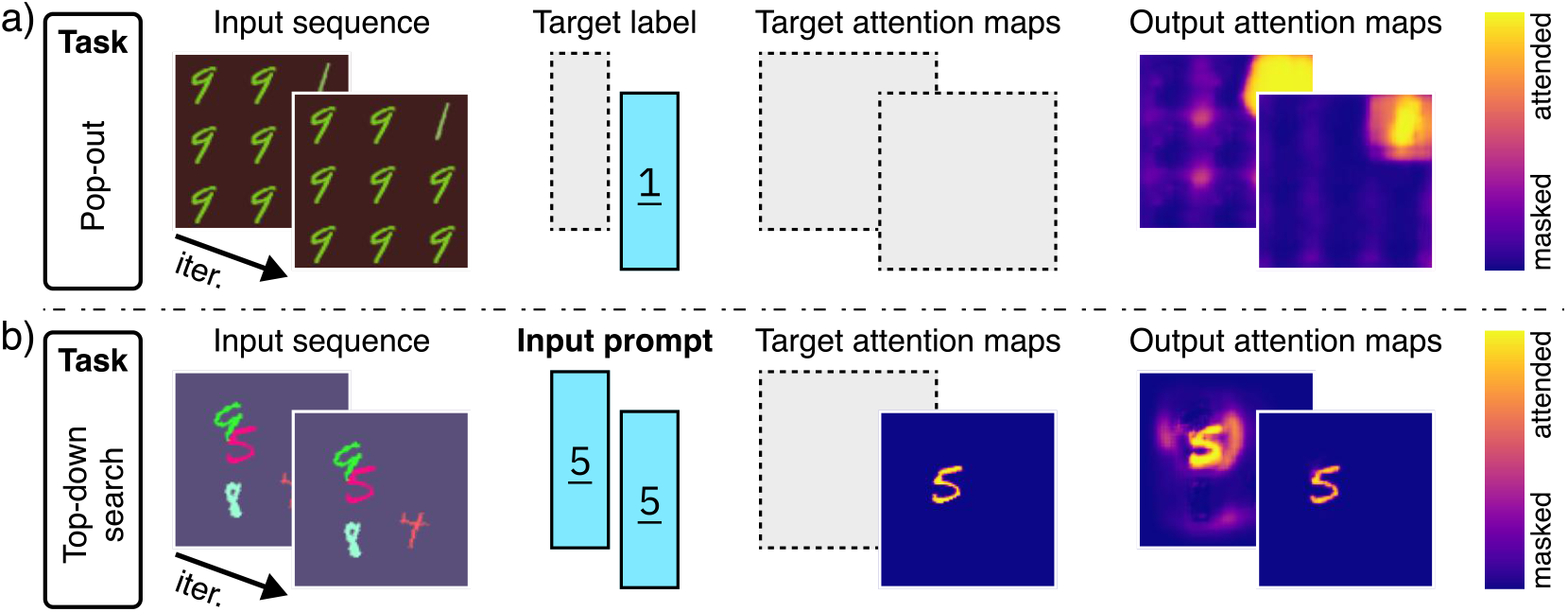
Multitask training on MNIST composites (Part 2/3). Results for a single model trained on seven tasks simultaneously. Here we present the results for: **a)** *pop-out saliency*, and **b** *top-down visual search*. For each task, from left to right: the input sequence, target labels or input prompts, target attention masks, and output (estimated) attention maps are shown. For the target label and target attention map, the gray dashed boxes imply that the target is NOT provided during training.

So far, attention has been guided by the input visual stimulus and mediated by the task input. However, it is often the case that we are given an explicit non-visual cue or description (i.e., prompt) of what to search for in a cluttered scene [Wolfe, 2010]. Since the prompt is the class or other distinct attributes of the object, the desired output of such tasks is the location of the prompted object. Despite the many kinds of visual search, here we focus on visual search through top-down attention. To demonstrate how our model can perform top-down search, we use an input composition of digits and an input class prompt that is fed to the bottleneck (i.e., top) (Fig. 3 b). The model is trained to find and attend to the prompted digit in the image. Our model achieves 88% attention accuracy and pixel error of 0.001 in recovering the correct attention map in the final iteration. Although here we restricted our task to the class, we have also successfully experimented with combination of other search attributes such as color (see supplementary materials).

A model of attention should be able to attend to an object as it moves through the scene. Spatial attention and awareness of the object’s location has been shown to be an important part of the binding process [Robertson, 2003]. Here, we will train the network on tracking and classify a moving digit (Fig. 4 a). The task is to fixate on and classify a stationary single digit for two iterations, and then to track the same digit as it begins to move through the scene with multiple distractor digits. Training for this task is fully supervised with target labels and masks provided during training, and achieves 99% classification accuracy and pixel error of 0.007 in recovering the correct attention map on the test dataset. Although the network was trained for only 7-iterations of tracking, it can track the target digit for a longer period of time during inference, indicating that the network can learn to track an object through space and time.

**Figure 4.**
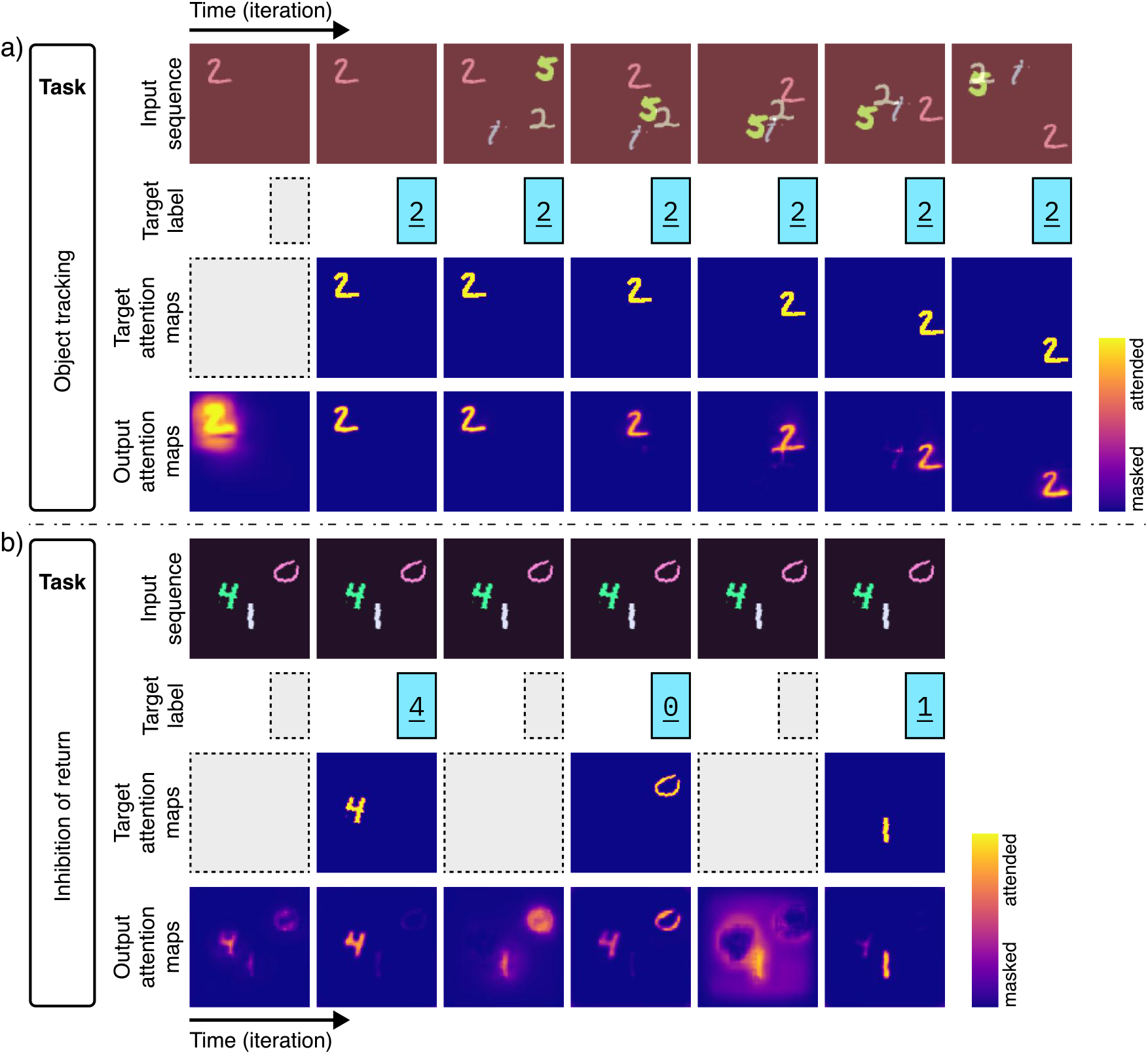
Multitask training on MNIST composites (Part 3/3). Results for a single model trained on seven tasks simultaneously. Here we present the results for: **a)** *spatial binding*, and **b)** *inhibition Of Return (IOR)*. For each task, from left to right: the input sequence, target labels, target attention masks, and output (estimated) attention maps are shown. For the target label and target attention map, the gray dashed boxes imply that the target is NOT provided during training.

When humans attend to multiple objects in a visual scene, they typically do so sequentially, moving from one object to the next [Klein and Ivanoff, 2008]. A large body of experimental evidence supports the idea of inhibition of return (IOR), whereby recently attended regions of a visual input are eventually inhibited, allowing the subject to move from one object to the next [Itti and Koch, 2001; Klein, 2000; Kwak and Egeth, 1992; Posner and Cohen, 1984]. Without inhibition of return, a subject will remain fixated on the most salient object. Here, we demonstrate that our model can learn to perform IOR on a multi-digit composite using only implicit training (Fig. 4 b). Our experiment is as follows: A sequence of identical compositions of *n* augmented digits from the MNIST dataset is fed into the model. The network learns to arbitrarily select, attend to and classify any of the digits in *k* given iterations. It must then move on to the next digit, and avoid returning to previously attended digits. We trained the network fully supervised simultaneously on *n* = 1, 2, and 3 objects with *k* = 2 iterations per digit. Our model achieves 96% accuracy in classifying all the digits in the scene and pixel error of 0.004 in recovering the correct attention for every digit. Thus, showing the network can readily cycle through objects in a multi-object scene.

#### Bregman’s illusion

Shared perceptual illusions between humans and artificial neural networks can inform us about the similarity of the underlying processes that are otherwise inaccessible to us [Benjamin et al., 2019]. Here, we want to investigate whether the model can account for well-known perceptual illusion for which its behavior is not explicitly predefined. Researchers have long known about the existence of Bregman perceptual illusion, in which a visible occlusion can provide cues that influence object perception. In the Bregman illusion, a stimulus such as a letter is occluded by an ink blott distractor (i.e., occlusion) (Fig. 5 a). When the distractor is removed, leaving an empty region, the stimulus appears fragmented and difficult to identify (Fig. 5 b). However, when the distractor is added, the fragments are contextualized as objects occluded by a foreground distractor, making the entire scene more interpretable (Fig. 5 a). The Bregman illusion is a classic demonstration of border ownership in Gestalt psychology and perception [Bregman, 2017; Qiu et al., 2007; Williford and Heydt, 2013].

**Figure 5.**
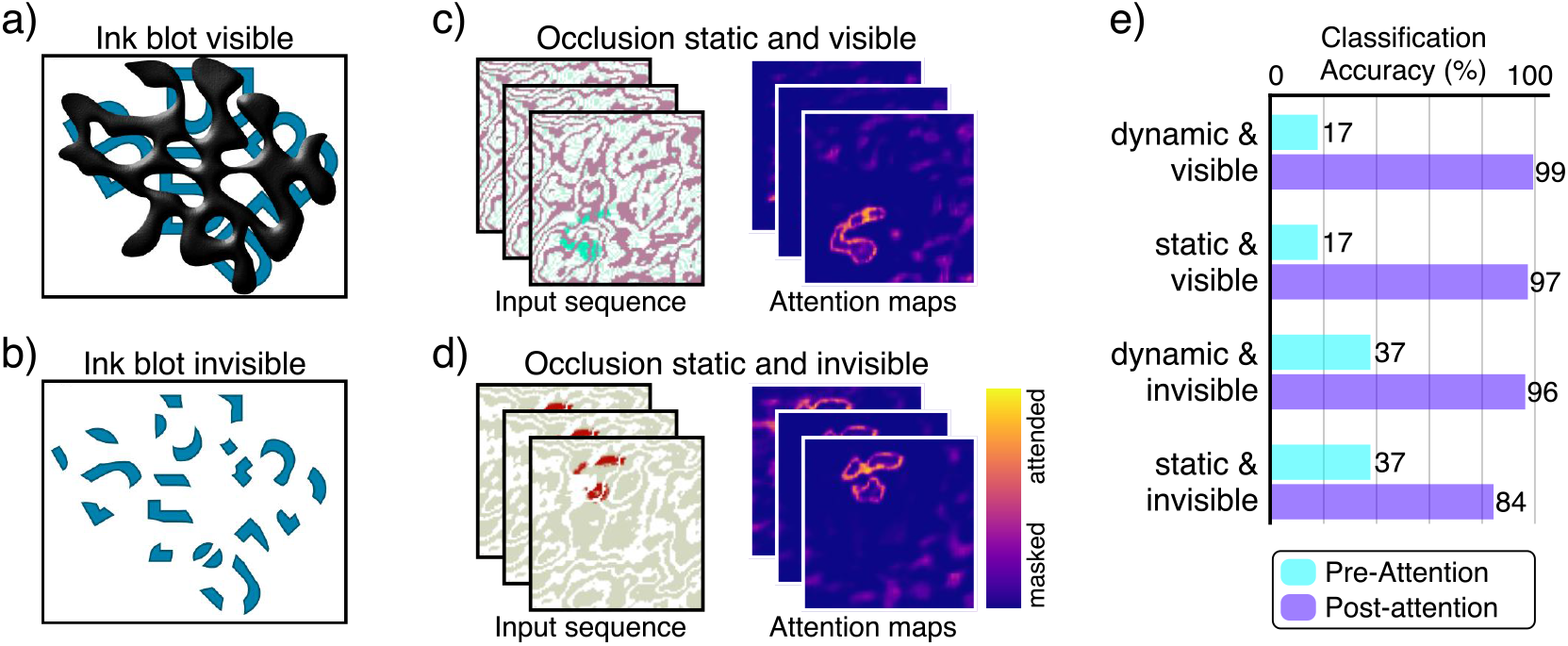
Bregman’s illusion. Bregman’s illusion is commonly used to demonstrate the concept of border ownership. a-b) Original illustration of the Bregman’s illusion, illustrating that visible ink blot helps with recognition of the letters. c-d) Similarly, visible occlusion in our experiment seems to help with recognition task and recovering the digit’s boundaries. e) Although visible occlusion appears to initially (i.e., pre-attention) hinder classification accuracy, attention seems to help the model to integrate occlusion boundaries to achieve higher performance.

To investigate this illusion, we start with the aforementioned model trained on the MNIST tasks. It is important to note that we did not explicitly train or fine-tune the model on this illusion, we only manipulate the stimuli used in the object recognition task (Fig. 5 c and d). In the original task, the foreground and background were moving and the model could slowly build a better representation of the object by integrating the information. Therefore we expect the classification and attention accuracy to suffer when the foreground is made stationary. The question is whether removing the foreground and its overlap with the target object would result in a change in classification and attention accuracy. Our analysis shows that the network does indeed perform better at recognition task when the occlusion is visible rather than when it is invisible, which is consistent with the experimental findings of Bregman’s illusion (Fig. 5 e). The results suggest that the presence of visible occlusion initially has a negative impact on classification, but the network has learned to use attention to extract boundary information from the occlusion and build a better representation of the digit (Fig. 5 e).

Although the MNIST dataset provides us with the freedom to design diverse tasks and evaluate the range of our model on multitask learning, the resulting compositions are far from those we encounter in nature. To further test our model on natural stimuli, we train a scaled up model on the animal subset of COCO [Lin et al., 2014]. We chose the COCO dataset because it contains fairly complex stimuli but also includes the true segmentation maps. Although the COCO dataset does not give us the flexibility of MNIST for multitask learning, we could devise three tasks to evaluate our model on: object recognition, top-down visual search, and perceptual grouping (Fig. 6). Since most images contain only a single animal or multiple animals of the same specie, we also trained on a composition of four distinct animals on natural background images from the BG-20k dataset [Li et al., 2022] for the top-down visual search (Fig. 6 c). For the object recognition task, we again used the BG-20k dataset, with a single animal randomly placed on the image, while the background moves slightly in each iteration (Fig. 6 a). Our model achieves 80% and 75% classification accuracy for object recognition and perceptual grouping tasks respectively. It is also effective in recovering the correct attention map with attention accuracies of 80%, 86%, 87% and pixel errors of 0.029, 0.029, 0.036 for for object recognition, perceptual grouping, and top-down search tasks respectively.

**Figure 6.**
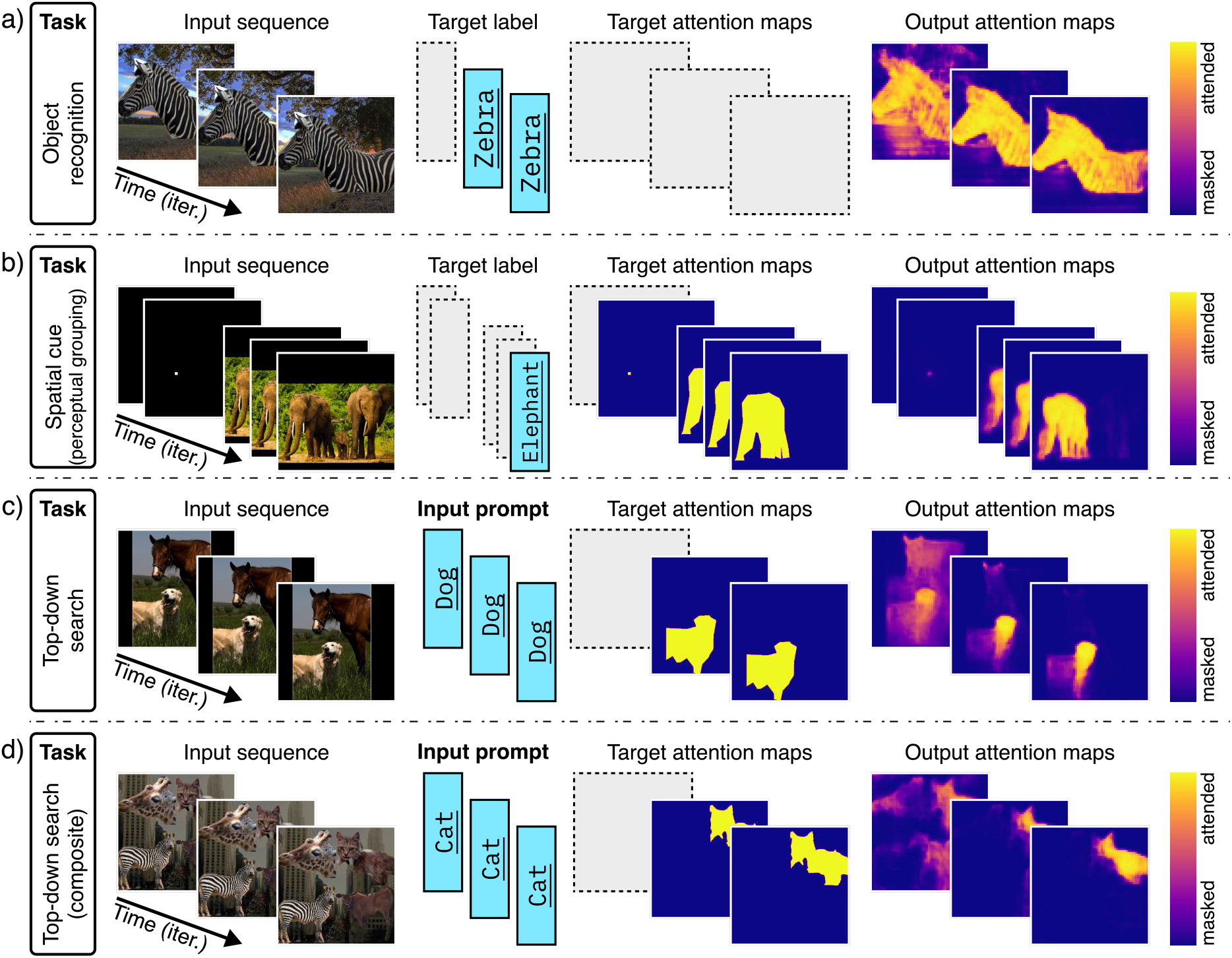
multitask training on COCO. Results for a single model trained on three tasks simultaneously: **a)** *object recognition*, **b** *cued perceptual grouping*, **c)** *top-down visual search*, and **d)** *top-down visual search on compositional images*. For each task, from left to right: the input sequence, target labels, target attention masks, output (estimated) attention maps, and classification and attention accuracy on the test data are shown. For the target label and target attention map, the gray dashed boxes imply that the target is NOT provided during training.

### Feature Attention and masking

One challenge in object recognition is the presence of spurious pre-attentive features in the scene. For example, most waterbirds are photographed near a lake or sea, which adds a blue background as a feature to the images. Studies in both human and artificial networks show that pre-attentive features like texture and color can lead to incorrect feature integration and misclassification [Geirhos et al., 2020; Geirhos et al., 2018; Lapuschkin et al., 2019]. Attention can help separate spurious and true correlations, by masking the irrelevant features and amplifying the informative attributes. To evaluate this theory, we use the CelebA dataset which contains images of celebrity faces and 40 binary attributes such as hair color and sex per image [Liu et al., 2015]. The images in the dataset are cropped such that the facial landmarks are approximately aligned across the dataset. Among the attributes in this dataset, there are some highly correlated features; namely, most celebrities with blonde hair color are labeled as female (Fig. 7 a). Therefore a naively trained neural network could use the blonde hair color as a low-level proxy for sex [Sagawa et al., 2019]. This makes the CelebA an interesting dataset to test whether our model can learn to attend to the right features for classification.

**Figure 7.**
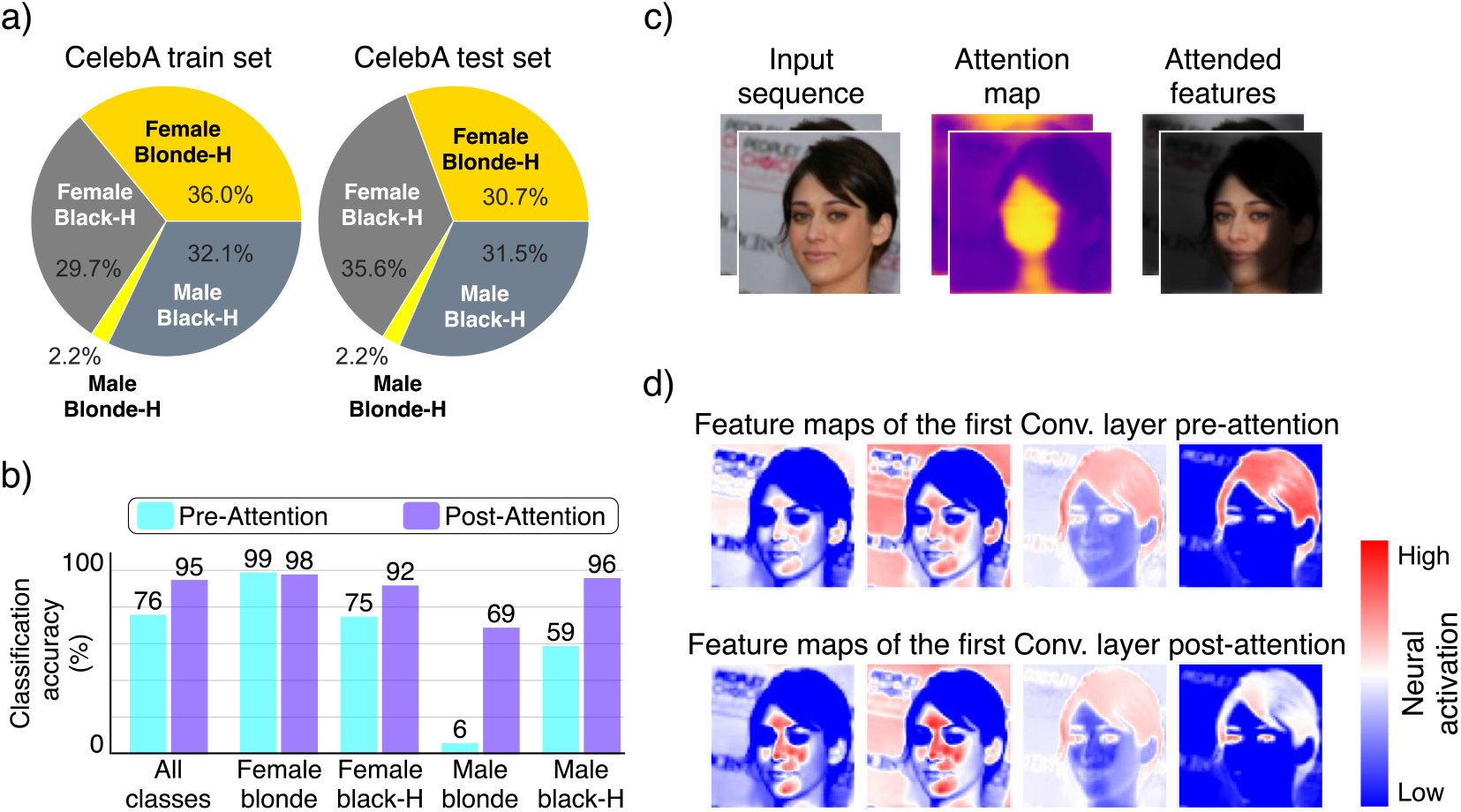
Feature Attention and masking. **a)** CelebA dataset contains a strong spurious correlation between hair color and sex. This spurious feature could be exploited by naive classifiers as a shortcut for sex. **b)** Classification accuracy per class and feature. Pre-attention results indicate that our model has not learned to correctly classify the expected sexes through the feature path alone, but rather uses recurrent attention to integrate the correct features. **c)** Purely trained on sex classification, our model has learned to attend to the facial features and ignore the hair and rest of the input. **d)** Feature maps in the forward path before and after attention.

To this end, we train our model on CelebA for a binary *sex* classification on sequences of length two, with NO target mask provided and without using class-weights to battle the class-imbalance (see supplementary materials for the extension of this task to localized attributes). The results suggest that the network does not learn to remove the hair information through the feature path, but rather uses attention to integrate the relevant features while masking out the hair to reach higher performance (Fig. 7 b, c and d). This could have important implications for how networks should learn to deal with spurious correlations, since removing the spurious features has been shown to be problematic [Khani and Liang, 2021].

### Neurophysiological Results

#### Object based attention

Inspired by the work of Roelfsema et al. [Roelfsema et al., 1998], we tested our model’s ability to perform object-based attention through curve-tracing task and examine the attention mediated neuromodulation in the network. Here, object is defined as a continuous and regular curve following gestalt principle of continuity. Curve-tracing is a task designed to investigate attentive neuromodulation, while reducing the influence of pre-attentive and overt processes. In the original curve-tracing experiment, two curves were drawn that might or might not intersect (i.e., switchbox). A cue was presented to determine the target curve and the subject was asked to covertly trace the curve. In our version of the curve-tracing experiment, we use contiguity as a cue for the model to decide which of the objects to attend to. The task is as follows: a cue is shown to the model for 2 iterations, followed by 3 iterations of two continuous, smooth, and equally salient curves randomly drawn from a third-order Bezier curve generator. One of the two curves is the target object, overlapping the previously shown cue, while the other curve is the distractor. Finally, the two curves disappear and two dots appear (training details are provided in the supplementary materials). The task is to attend to the dot that overlaps with the target object (Fig. 8).

**Figure 8.**
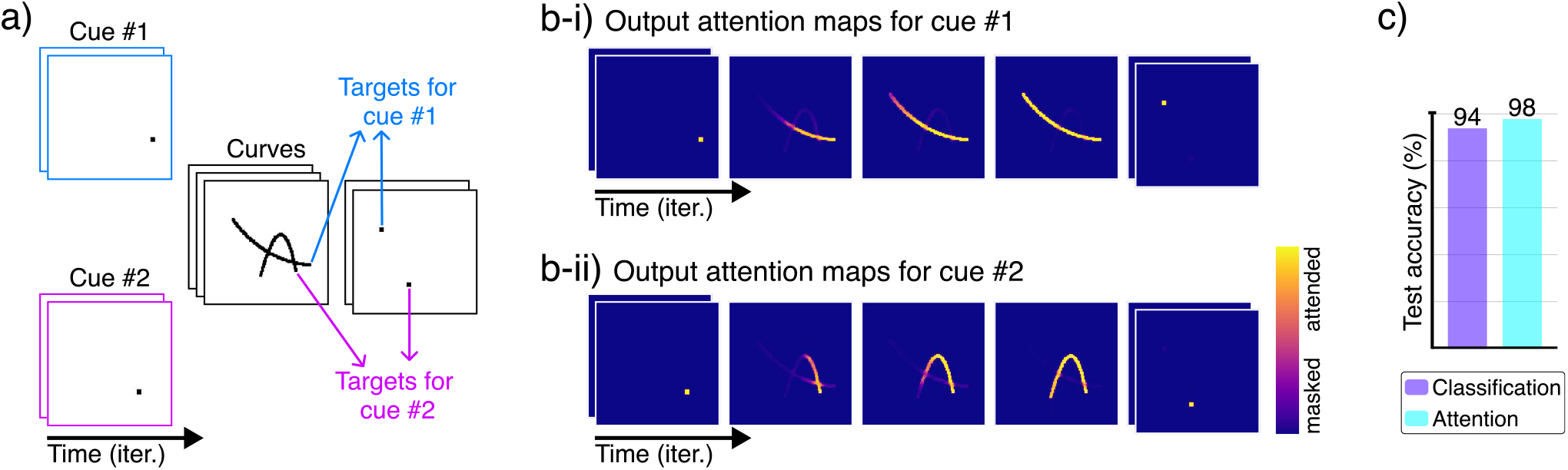
Object-based attention. **a)** The input sequence for the curve-tracing experiment consists of 3 phases from left to right: i) fixation (i.e., visual cueing) for 2 iterations; ii) stimulus (i.e., curve tracing) for 3 iterations; iii) saccade (i.e., attending to the target decision point) for 2 iterations. For any two curves and decision points, we can use two distinct cues to dictate which curve and decision point are the targets. **b-i)** and **b-ii)**) estimated attention maps for two sequences that share the same curves and decision points but different cues shown in the left-side of this figure (i.e., a). **c)** Classification accuracy for the correct decision point and attention map accuracy for the target curve.

An advantage of this experiment is that for any combination of two curves, we can use the initial cue to switch the target and distractor curves, giving us the control we require for the experiment. Not only is our model able to learn and perform the task correctly; i.e., it can learn contiguity, continuity, and regularity to attend to the correct sequence of stimuli, but also its neural activity is consistent with the findings from the primary visual cortex of the macaque monkey performing a similar experiment [Roelfsema et al., 1998] (Fig. 9). We can show that attention positively modulates the activity of neurons with the target curve in their receptive field, compared to when the same object is the distractor (i.e. unattended).

**Figure 9.**
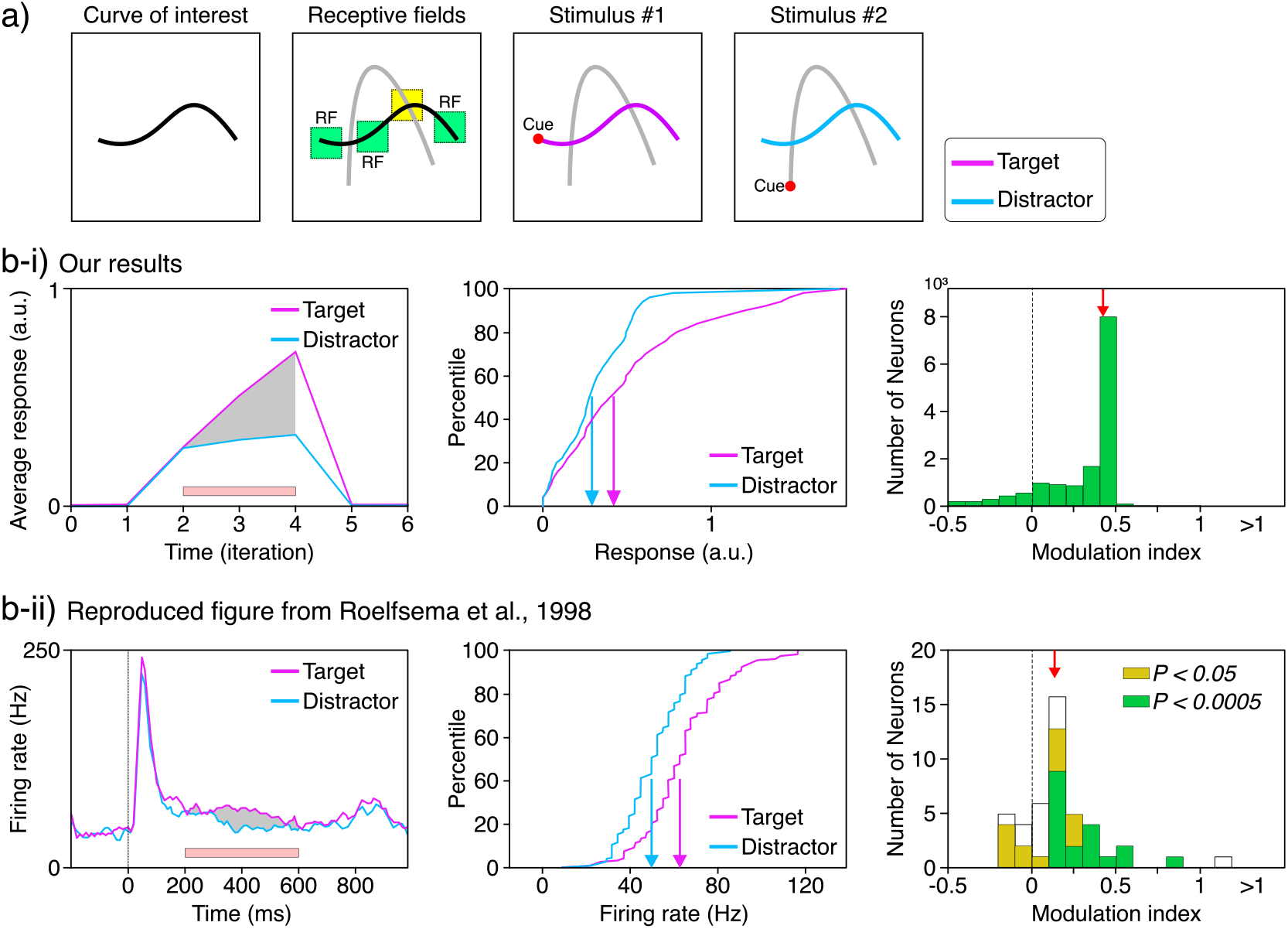
Attentional Neuromodulation. **a)** For any two curves, we find all the neurons that their receptive fields include one curve but not the other (here, green squares are considered acceptable receptive fields while the yellow one is an example of rejected receptive field). We can then use the visual cue to mark any curve as target object (stimulus #1) or as distractor object (stimulus #2). **b)** Activity of neurons is enhanced through attention. Here we compare the results of **b-i)** neurons in first layer of our model for the curve-tracing experiment with **b-ii)** results from V1 neurons in macaque monkeys performing a similar task [Roelfsema et al., 1998]. The left graphs show neural response through time for the target (i.e., attended) versus the distractor (i.e., masked) curves. Our plot shows the average response for all the selected neurons in the first layer. The pink bar marks the curve tracing window, used for the statistical analysis. The middle graphs demonstrate the response distributions of neurons for the target and distractor scenarios. The right graphs depict the histogram of response modulation index, showing significant positive modulation of neuronal activity. The red arrow indicates the median modulation index. The graphs are made in similar style to those in [Roelfsema et al., 1998].

#### Attention-invariant tuning

Tuning curves are often used to depict how a neuron changes its activity (i.e., firing rate) in response to a change in a single attribute of its input. A common tuning curve for neurons in early visual areas which are sensitive to stimuli similar to 2-dimensional Gabor wavelets, is the orientation tuning. To extract the tuning curves, the experiments are designed so that the neuron’s activity is mostly driven by input from early visual areas and is free of recurrent or higher-level signals. However, here we are interested in how the neuron changes its tuning in the presence of attention. Neurophysiological studies suggest that when a subject is attending to an object, neurons that are associated with the object’s receptive field show a stronger activation than when the subject is not attending to it [Lee and Maunsell, 2010; Maunsell, 2015; McAdams and Maunsell, 1999; Treue and Trujillo, 1999]. Furthermore, [McAdams and Maunsell, 1999] showed that the change in the activation is multiplicative, meaning that other attributes such as width and preferred orientation are statistically unaffected, suggesting that neural tuning is attention-invariant. Like biological neurons, neurons in the early layers of an artificial neural network trained on naturalistic images exhibit tuning curves that resemble Gabor wavelets. In the deeper layers, however, the learned filters become increasingly complex, resembling the tuning curves of neurons in the ventral visual pathway (Fig. 10 a). Here we aim to test whether the neurons in our model also exhibit attention-invariant tuning.

**Figure 10.**
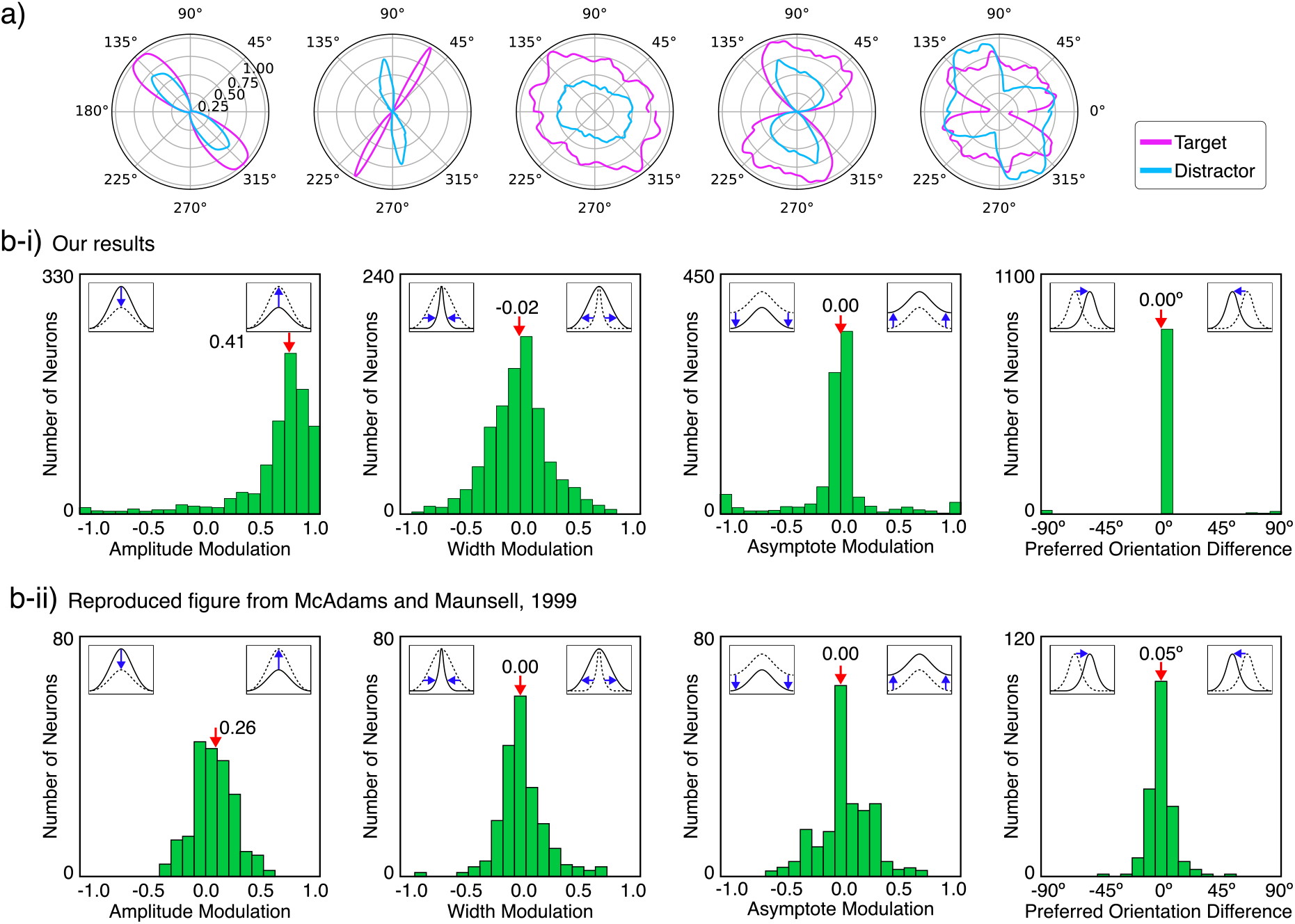
Attention-invariant tuning. **a)** Orientation tuning curves of five neurons in different layers of our model trained on the curve-tracing task. For the analysis of the attention-invariant turning, we only consider neurons with Gaussian-like tuning curves (e.g. the two plots on the left) and reject the rest. **b)** Attention increases the response amplitude but has little to no effect on the width, asymptote, and preferred orientation of the tuning curves. **b-i)** our results from the penultimate feature layer in our model compared to **b-ii)** the findings from V4 cortical neurons in macaque monkeys [McAdams and Maunsell, 1999]. The red arrows and values show the median. The graphs are made in similar style to those in [McAdams and Maunsell, 1999].

We start with the model trained on the curve-tracing task and follow the method used by [McAdams and Maunsell, 1999]. As input, we use the composition of two distant bars (i.e., short straight lines). We use visual cues to switch attention between the two bars, creating two states of attended and unattended, for the target bar. We then extract neurons that have the target bar in their receptive field. Furthermore, we will only consider those neurons that have a bell-curve (i.e., Gaussian like) orientation tuning. Finally, we can show that attention affects the neural activation in later layers of our network through multiplicative scaling without changing the overall shape of their tuning curves, similar to experimental findings from [McAdams and Maunsell, 1999] in V4 cortical neurons (Fig. 10 b).

## Discussion

We have demonstrated that a substantial portion of our understanding of visual attention can, in principle, be explained by bidirectional recurrent gating mechanism. We have introduced a unifying model that integrates multiple attentional phenomena observed in the cognitive and neural sciences. Our model performs similar to humans, both qualitatively and quantitatively, on a range of canonical visual tasks, including object recognition, top-down search, pop-out effects, inhibition of return, and visual inference. Additionally, it replicates various neurophysiological findings, such as attention-mediated neuromodulation and attention-invariant tuning. Computationally, we showed how our model can correctly mask spurious features in the images, leading to more robust predictions. This work shows that bidirectional recurrent gating is a powerful mechanism for modeling and understanding attention.

Our model, naturally, has its limitations in describing neural and behavioral effects. For example, it does not account for overt attention mechanisms such as saccades and foveation, which are crucial for visual attention in humans [Itti et al., 2005]. Fortunately, effective models of these mechanisms exist and could, in principle, be integrated into our framework [Mnih et al., 2014]. Additionally, our model does not explicitly address aspects of neural computation such as non-stationarity, time-varying dynamics, and oscillations (beyond cycling through objects). However, given the recurrent nature of our network, we believe it can be adapted to incorporate these dynamics. Clearly, while our model describes a range of effects, there are still many findings it cannot yet quantitatively account for.

Next, our model uses Backpropagation Through Time (BPTT) to update its weights. While BPTT is a powerful tool in machine learning, there is no direct evidence that backpropagation is used as an updating mechanism in the brain [Lillicrap et al., 2020], especially not in a BPTT setting where information would need to travel backward in time. Nevertheless, we argue that our mechanism provides a key inductive bias enabling the network to perform well on multiple attention tasks, although it currently relies on BPTT. However, computationally, all we need is gradient descent and many approximations may exist. Biologically plausible approximations of gradient descent have extensively been studied [Bengio et al., 2015; Richards and Kording, 2023]. In fact, backpropagation has been used to learn brain-like representations for visual tasks [Cadena et al., 2019; Khaligh-Razavi and Kriegeskorte, 2014; Kriegeskorte et al., 2008; Kubilius et al., 2019; Rajalingham et al., 2018]. Similarly, while weight sharing in convolutional layers has limited biological plausibility, despite being inspired by simple and complex cells in the visual cortex [Fukushima, 1988]. Recent models have proposed augmentations or wake-sleep cycles to train networks resembling emergent weight sharing [Pogodin et al., 2021]. In other words, our biologically motivated mechanism facilitates modeling attention in artificial neural networks through backpropagation, and we do not consider the biological plausibility of deep learning methods to be a limitation of the mechanism.

We have demonstrated that the proposed multitask learning paradigm can reduce the necessity of full supervision in some tasks. However, so far, our system still requires supervised signals. How do humans obtain supervised information about segmentation? Arguably, they do so by observing moving objects. The movement of these objects effectively separates them from the background, providing a signal that could be considered largely equivalent to supervised segmentation information. Similarly, additional strong cues exist for other aspects we use in supervised learning. For instance, abstract knowledge about object identity is often encoded across different modalities; for example, the presence of a horse is associated with a distinct smell. While our approach relies on supervised signals, we maintain that exploring what cues and learning algorithms the brain actually employs is a separate, albeit relevant and intriguing, research question.

Despite the drawbacks, our model of object-based attention reasonably explains data with the ability to solve real-world problems, and can efficiently learn to perform complex behavioral tasks. More importantly, we realized this with a scale-able, modular, and stable architecture that is built upon basic neural network components. Although marginally requiring more computation compared to feedforward convolutional neural networks, attention and recurrency are far too important to ignore in models of visual cognition [Lindsay and Miller, 2018]. Therefore, we argue that our model would provide a pragmatic and effective alternative to existing CNN architectures commonly used to model the visual system [Lindsay, 2021; Xu and Vaziri-Pashkam, 2021]. This would also enable design of more complex experiments that are closer to real-world behavior [Matusz et al., 2019].

Our network brings recurrency and gating, two fundamental computation schemes ubiquitous in cortex, to better model the biological visual system. Combined with other elements like hierarchical feature representation learning, the system has the right structure to model neurophysiological data. Furthermore, the multitask paradigm opens the door to design concurrent tasks that could bring about emergent behaviors at both behavioral and neurophysiological levels.

Importantly, our model provides a sufficiently complex baseline model that can be used to develop better models of object-based attention in the brain. Our results state the positive impact of attention and multitask learning on generalization and robustness [Cheung et al., 2019; Driscoll et al., 2024; Sinz et al., 2019; Thorat et al., 2019]. Our model of object-based attention motivates new insights into our understanding of decision-making in a broad range of visual tasks, including credit assignments, binding, and memorability. This promises to be a starting point for further research in vision, learning, and binding that combines real-world tasks with cognitive modeling. Simultaneously, with the recent rise of interest in object-based attention and binding in the field of machine learning [Greff et al., 2020; Locatello et al., 2020], we see our network as a productive common ground between neuroscience and machine learning.

We also propose that the deep learning literature may benefit from incorporating insights from such attention modeling. By introducing attention as a primitive in the standard artificial neural network framework, we provide a flexible mechanism that allows stimuli, prompts, tasks, and outputs to jointly determine what is attended to and propagate this information through a large network. This approach fundamentally acknowledges the compositionality of the real world. Further research is needed to identify the types of tasks where incorporating human-like attention enhances network performance.

## Acknowledgements

This work was partly funded by the German Ministry for Education and Research (under refs 01IS14013A-E, 01GQ1115, 01GQ0850, 01IS18056A, 01IS18025A and 01IS18037A) and the German Research Foundation (DFG) within the RTG 2433 DAEDALUS. Furthermore, KRM was partly supported by the Institute of Information & Communications Technology Planning & Evaluation (IITP) grants funded by the Korea government (MSIT) (No. 2019-0-00079, Artificial Intelligence Graduate School Program, Korea University and No. 2022-0-00984, Development of Artificial Intelligence Technology for Personalized Plug-and-Play Explanation and Verification of Explanation).

## Code and data availability

The source code is available on Github: https://github.com/ssnio/bio-attention or upon request. The datasets are publicly available on their corresponding online repositories. The scripts to generate the composites are part of the source code.

## Supplements / Appendix

### Validation

We followed the DOME recommendations and framework presented by [Walsh et al., 2021] with respect to assessment and reproducibility.

#### Data

The data samples are task-specific and are generated by augmenting and composing images from publicly available datasets: MNIST [LeCun, 1998], COCO [Lin et al., 2014], and CelebA [Liu et al., 2015]. We use different image augmentation (such as random flip and rotation) to improve the generalization. Regarding the train-test splits, the CelebA dataset comes with three distinct sets: training, validation, and test datasets that we use accordingly. But the MNIST and COCO datasets do not include a dedicated publicly available and labeled test dataset, so we use cross validation on the train set and use the validation set for testing. In all experiments, the validation sets are only used for inference to find the best trained model and the test dataset is used for final reporting and visualizations. The data samples for the curve-tracing experiment were created randomly using Bezier curves.

#### Optimization

All models are trained on a single NVIDIA A100 GPU. We use Adam algorithm as the gradient descent optimizer with weight decay regularization. We perform grid search for finding a learning rate and regularization factor that yields better validation error. We found that very small learning rates often lead to better results (i.e., more stable and more reasonable attention maps). Our loss function could be written as the composition of three weighted objectives:

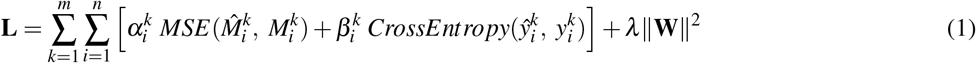

where subscript *i* is the iteration and superscript *k* denote the task index. The first term inside the sum is the Mean-Squared Error (MSE) loss for the estimated 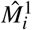 and true attention maps *M*_*i*_, if available. Next is cross-entropy as the classification loss of output labels. For a given experiment, we control for the existence and importance of each term in the iteration *i* with the hyper-weights *α*_*i*_ and *β*_*i*_. The last term is the *L*_2_ regularization term for the learnable weights **W**. For the COCO model, we also used class-weights to improve the performance.

**Table 1.** Training hyper-parameters.

| | n-Epochs | Batch-size | Learning rate $\eta$ | $\eta$ Scheduler milestones | scheduler $\gamma$ | L2-rate $\lambda$ |
| --- | --- | --- | --- | --- | --- | --- |
| MNIST | 96 | 128 | $5 \times 10^{-4}$ | [32, 64] | 0.2 | $1 \times 10^{-6}$ |
| COCO | 42 | 128 | $5 \times 10^{-4}$ | [24, 32] | 0.2 | $1 \times 10^{-5}$ |
| CelebA | 32 | 128 | $2 \times 10^{-4}$ | [16, ] | 0.1 | $5 \times 10^{-4}$ |
| Curve Tracing | 64 | 128 | $5 \times 10^{-4}$ | - | - | $1 \times 10^{-6}$ |

#### Model

The network is built by stacking multiple modules of bidirectional recurrent gating blocks that could have different “processing and learning layers” (Fig. 11). The first iteration always starts with a flat attention map. As the image passes through the feature path, the model extract higher level features and builds a richer representation of the input. The model also stores the feature maps for each layer to be used later by the attention path. Finally, the feature path ends (if necessary) with dense RNN layers that act as working memory. The output of the last layer in forward path is used for label classification. In most experiments, we do not consider the first label prediction for classification error since the first feature pass is unaware of the task and is the results of flat attention. The attention path starts with the output of the forward path concatenated with the task encoding. Note that for the top-down tasks, the forward output is discarded and the one-hot encoded prompt is fed into model instead. After attention RNN layers, the resulted activity is concatenated with the feature maps from the corresponding forward layer and scaled according to the task-embedding. This signal now contains information about the task, the feature representations, and latent information and can be used by the consequent layers (namely transposed-convolution, normalization, and non-linearity) to generate the attention maps for corresponding forward layer. Other than the bottleneck, the concatenated features and attention maps, form a tensor of shape (channels, height, width). The task embedding modulates the channels and not in the pixel level. This process is repeated for every layer in the attention path until the attention map reaches the bottom of the model, which concludes the first iteration. The next iterations is similar to the first iteration except the inputs to each layer are now modulated by the scaled attention maps from the previous iteration. We use a multiplicative scale of 0.5 to allow partial attention and avoid losing all the distractor information, as they might be relevant for later iterations [Maunsell, 2015].

**Figure 11.**
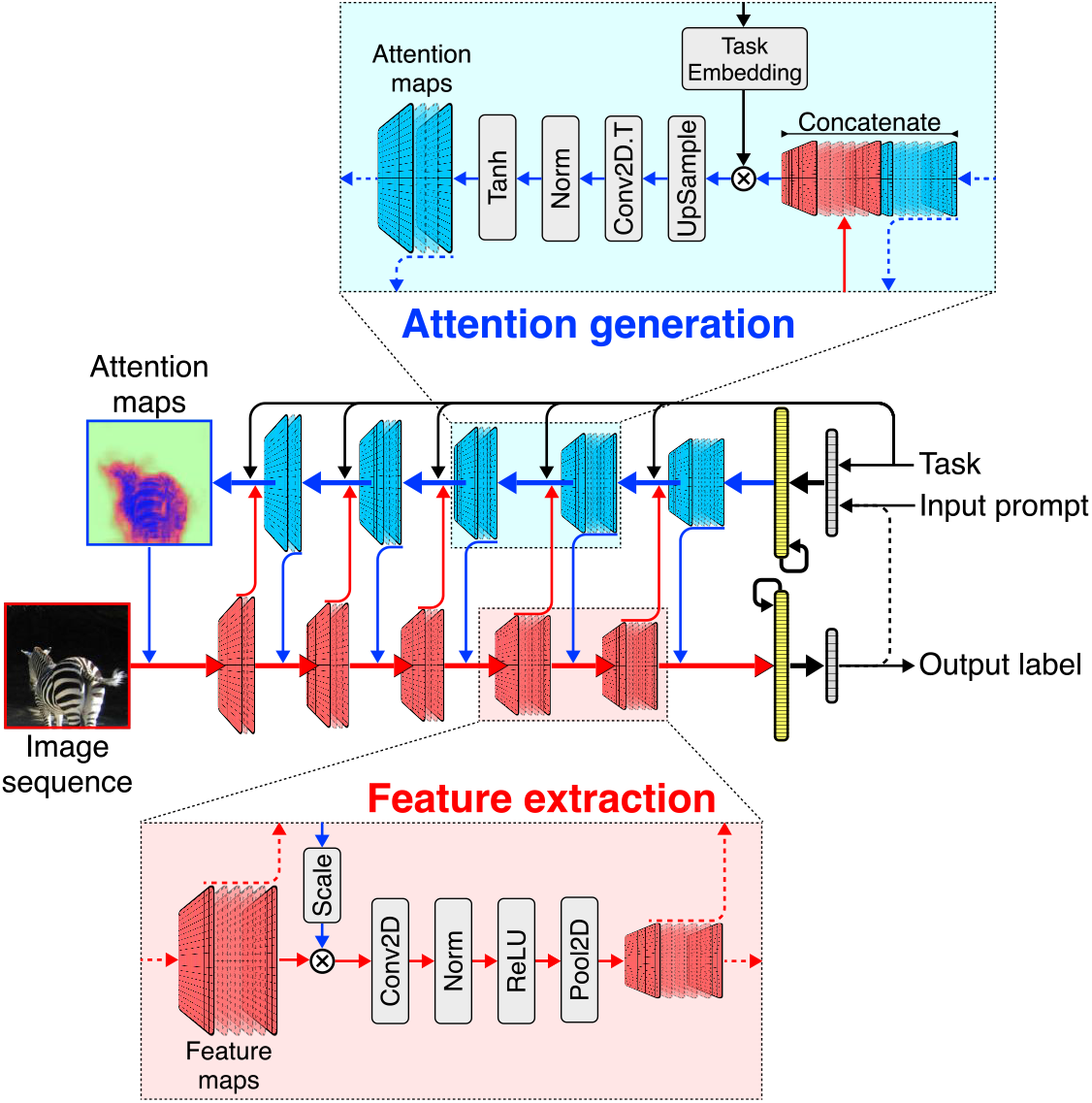
Architecture schematic. The elements of the two pathways are convolution (Conv2D) and transposed-convolution (Conv2D.T), nonlinear activation functions (ReLU and Tanh), pooling and up-sampling, and divisive normalization (Norm). The feature map 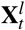 of block *l* at iteration *t* is multiplicatively modulated by the attention map 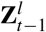 from the previous iteration before being processed through the forward block *l* + 1. The output of block *l* + 1 is passed to the corresponding block in the attention path, concatenated with the attention map *l* + 1 before modulation by the task-embedding. The attention block then creates the attention map 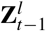 for the next iteration

#### Model details

Although all the models used are similar in their core architecture, each network is scaled and equipped differently depending on the task and data complexity. We mostly tried to only keep elements that are crucial for the experiment. For example, in the MNIST model we used RNN layers as working memory which is necessary for the IOR task (Fig. 12). For COCO we used a scaled up version of MNIST model, but did not include the RNN layers. We also use GELU (Gaussian Error Linear Unit) activation function for COCO model, since it has been shown to improve training [Hendrycks and Gimpel, 2016]. Moreover, since the COCO model was trained on only three tasks, we found that it does not require task-input in all the decoding layers (Fig. 13). A common theme between all the models is a bottleneck that is wider than the number of classes. We choose a wider bottleneck to allow non-class features that might be relevant for solving the tasks to flow to the attention path. This seemed to improve the network performance.

**Figure 12.**
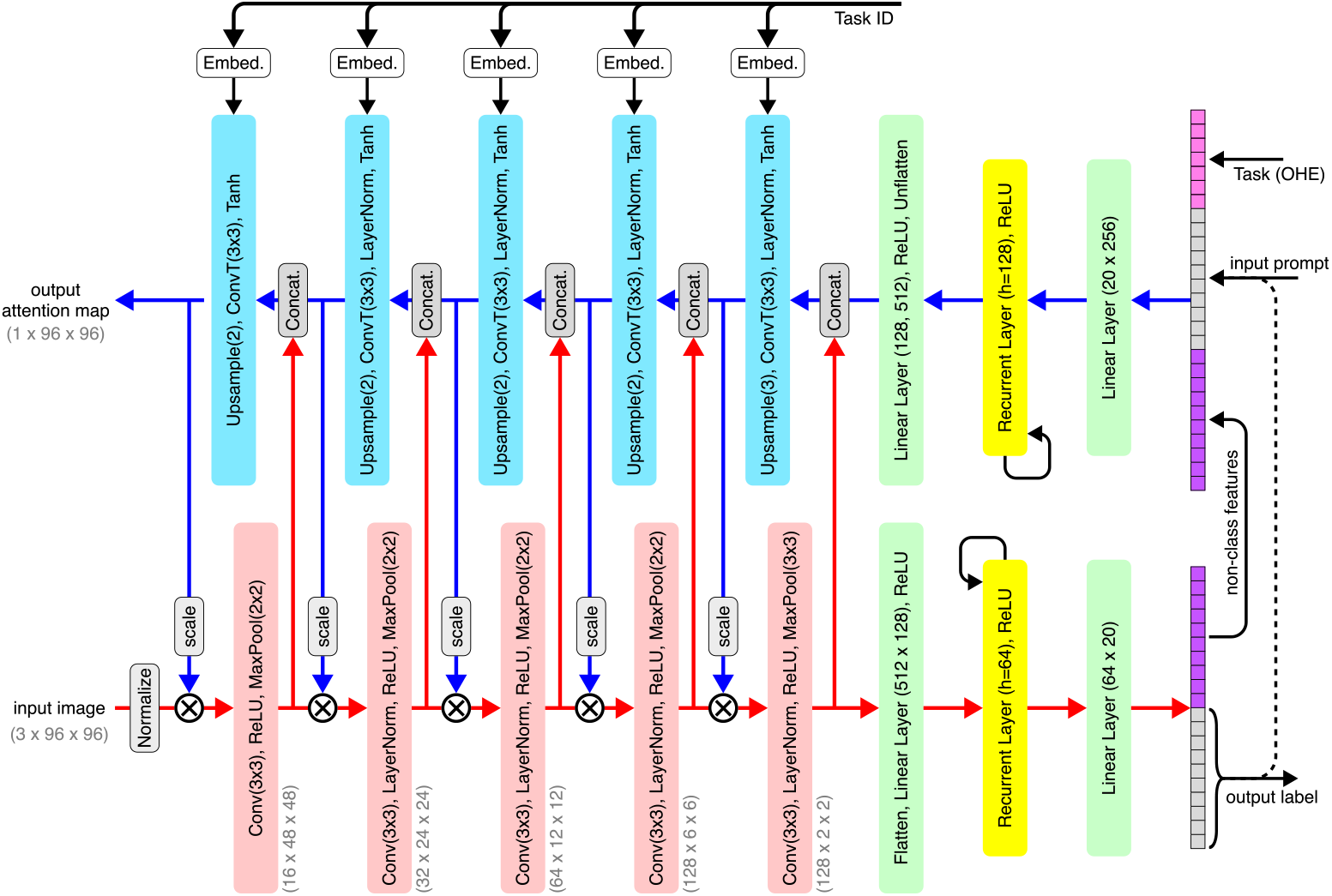
Detailed architecture used for the MNIST experience.

**Figure 13.**
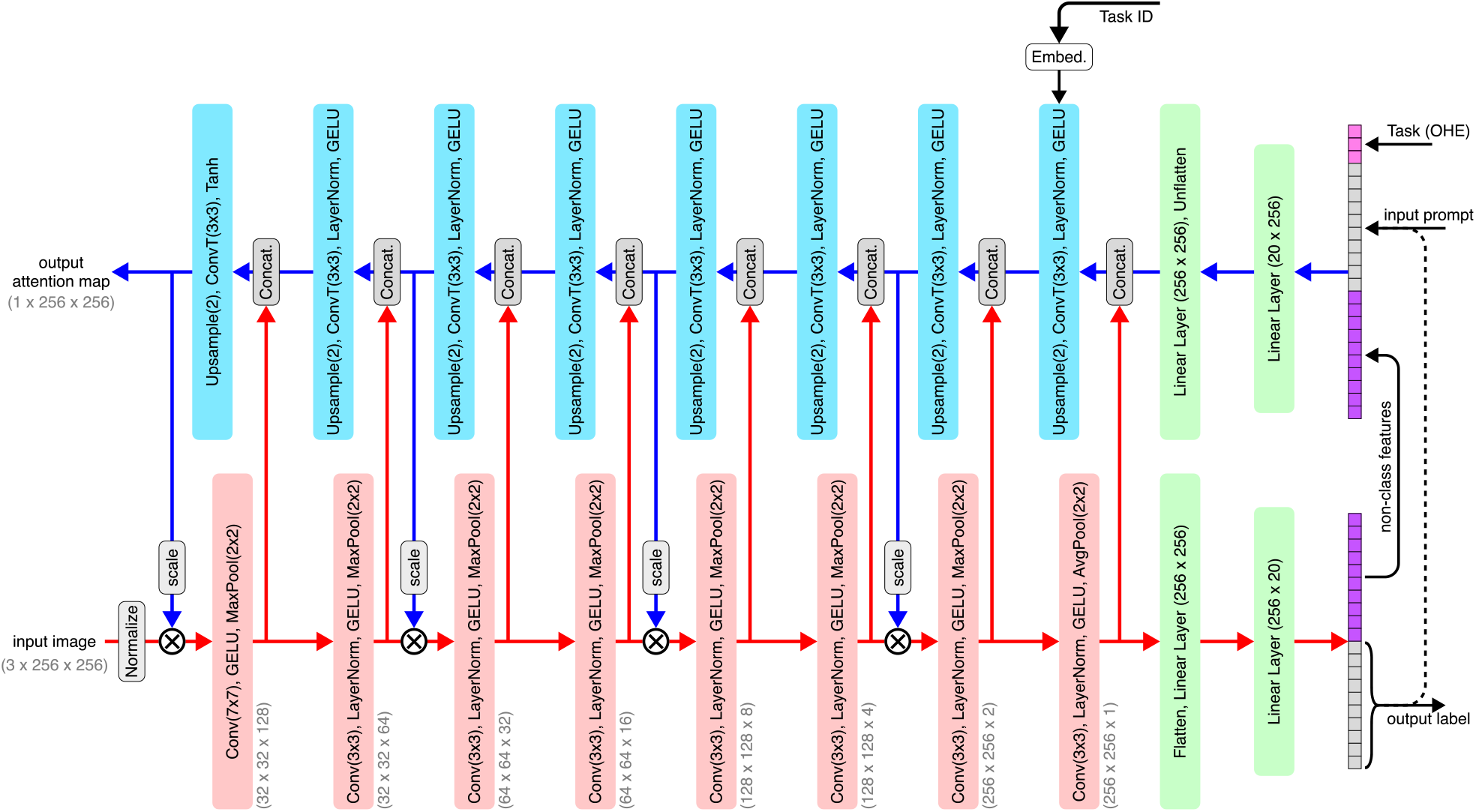
Detailed architecture used for the COCO experience.

#### Evaluation

The code and trained models are all made available for validation and further investigation (please see Code Availability section). For classification, we use Cross-Entropy loss. The MNIST dataset is sufficiently balanced, but for COCO we used class-weights to combat the imbalanced class distributions. For CelebA experiment, although the dataset is not balanced, we did not use class-weights to rather emphasize on the networks ability in learning without prior knowledge of class distribution and spurious correlations.

For segmentation, we use mean-squared loss for training, but we use attention accuracy and pixel error for reporting as they are more intuitive and easier to interpret.

### Attention accuracy

Attention accuracy measures the normalized accuracy of a pixel being correctly attended or masked (i.e., the target and estimated pixel value having the same sign). We use normalization to correct the target to scene area ratio, leading to chance level to always be at 50% (i.e., the whole scene being attended or the whole scene being masked). The measure is similar to the Intersection over Union (IoU) commonly used in computer vision [Rezatofighi et al., 2019]. The following equation explains the measure:

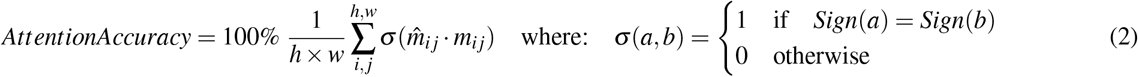

where 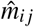 and *m*_*i j*_ denote the *i j* − *th* pixel of estimated and true attention maps respectively, while *h* and *w* are the height and width of the attention map.

**Pixel Error** Pixel Error denotes the mean squared euclidean distance between estimated and true attention maps.

**Modulation index** We use the Modulation Index (MI) similar to those described in [Roelfsema et al., 1998] and [McAdams and Maunsell, 1999]:

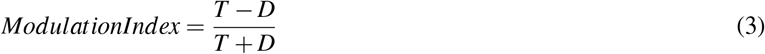

where *T* and *D* denote the averaged response (i.e. activity) of neurons to the target and distractor curves respectively. We do not use factor 2 and resulting in MI that is always in [−1, 1] range.

### Bell-curve fitting

For the invariant tuning experiment, we used the following method to accept or reject a tuning curve based on their shape (i.e., whether they are bell-liked). Starting with the raw tuning curve *y* with length 180 (i.e., domain of 0 to 179 degrees), we first use a 1-D Guassian filter to smooth out the tuning curve. The following class calculates the curve attributes (namely: amplitude, asymptote, width, and preferred orientation) and returns the mean-euclidean distance between the tuning curve and a bell-curve that has similar attributes.

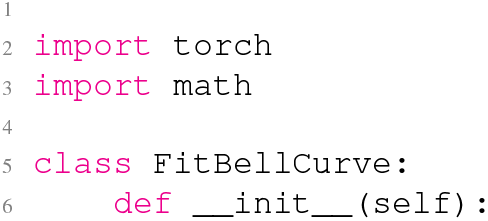
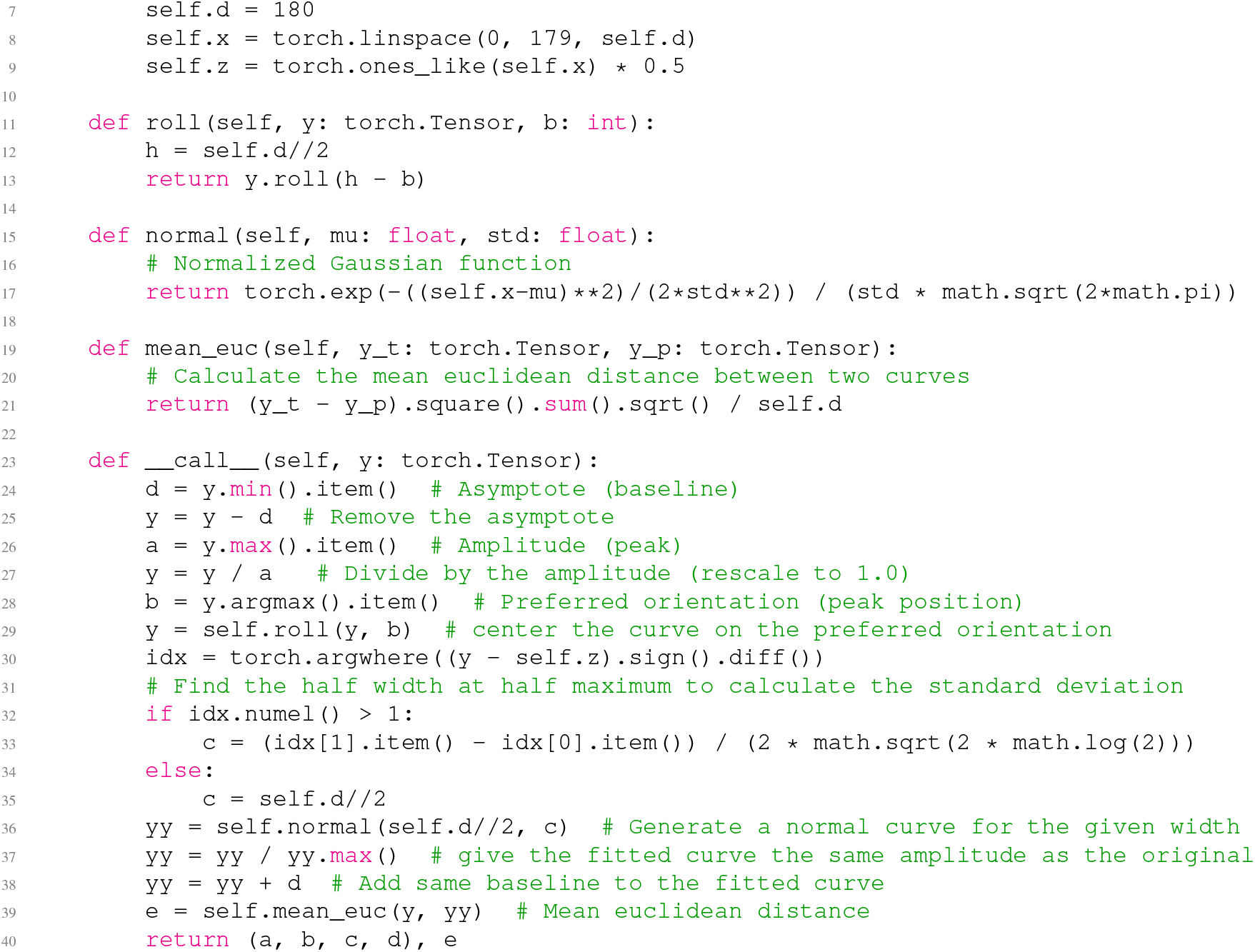

### Classification tasks for CelebA attributes

A challenge in sex classification is the many correlated features that are present in the CelebA dataset. For example, wearing eyeglasses is strongly correlated with the male class, while having blonde hair is more common between the female celebrities in the dataset. Here, we look at features that are localized and hence the learned attention maps are easier to interpret. We train separate models solely on binary classification of: *sex* (Fig. 14 a), whether the person is *smiling* (Fig. 14 b), whether the person has *blonde or black hair* (Fig. 14 c), and whether the person is *wearing eyeglasses* (Fig. 14 d). The results show that the network trained on smiling puts more attention to the mouth, lower cheeks, and chin (Fig. 14 b), while the network trained on eyeglasses seems to attend strongly to the mid-face and eyes (Fig. 14 d), Surprisingly, the model trained on hair color seems to attend not only to the hair but also the eyes (Fig. 14 c).

**Figure 14.**
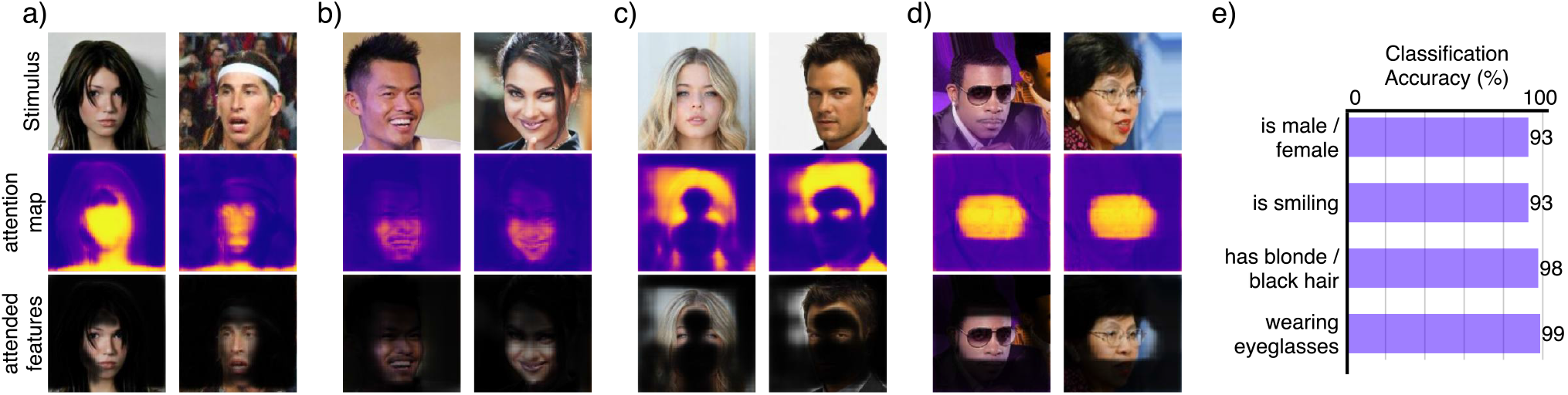
Binary classification for localized face attributes. Input image sequence, output attention map, and attended features for different models trained on binary classification of **a)** sex, **b)** smiling, **c)** hair-color, **d)** and eye-glasses.

### Curve-tracing task

Training of the curve-tracing task is done entirely through the target attention maps (i.e., no classification error) (Fig. 15). We could successfully train the model to perform the task, with and without target curve supervision. The results shown in the main text are from the model that received target curve supervision (Fig. 15 a).

**Figure 15.**
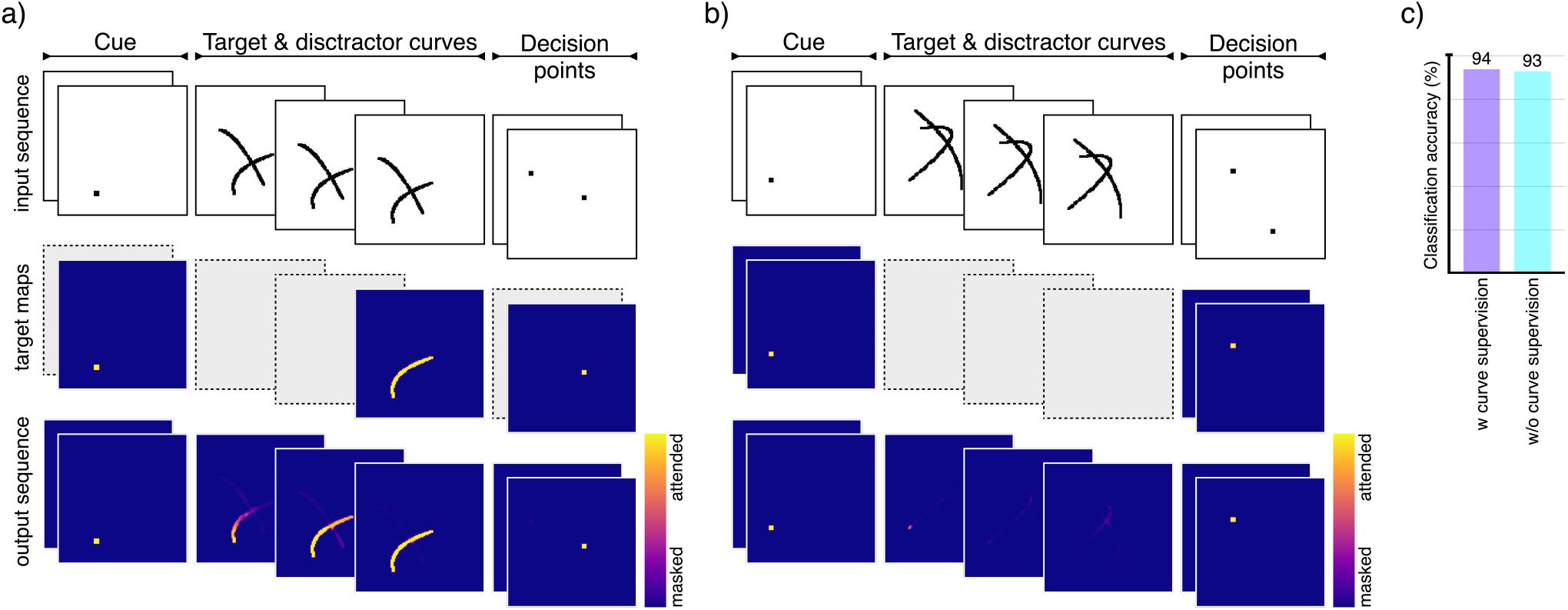
Training approach for the curve-tracing task. **a)** Full supervision: Target curve is provided while training on the task alongside the cue and target decision point. **b)** Partial supervision: Only the target attention maps for the cue and target decision are provided during the training, so the network has to learn about the curves and their abjectness by itself. The gray dashed boxes imply what is NOT provided during training. **c)** Our model can learn to perform the task through either of the training regimes, although the partial supervision took longer to train.

There are some interesting observations to be made from this experiment. For example, the model seems to have an inherent attention (or grouping) propagation behavior (Fig. 16 a versus b) which could be further investigated to see if our model produces similar results to those from [Ekman et al., 2020] on top-down object-based attentional spread. Also, even though the model is really good in separating the two curves in most scenarios (e.g., Fig. 16 c), it fails to make a decision (i.e., choosing a single point) when the curves are ambiguously placed, and attends to both decision points (e.g., Fig. 16 d).

**Figure 16.**
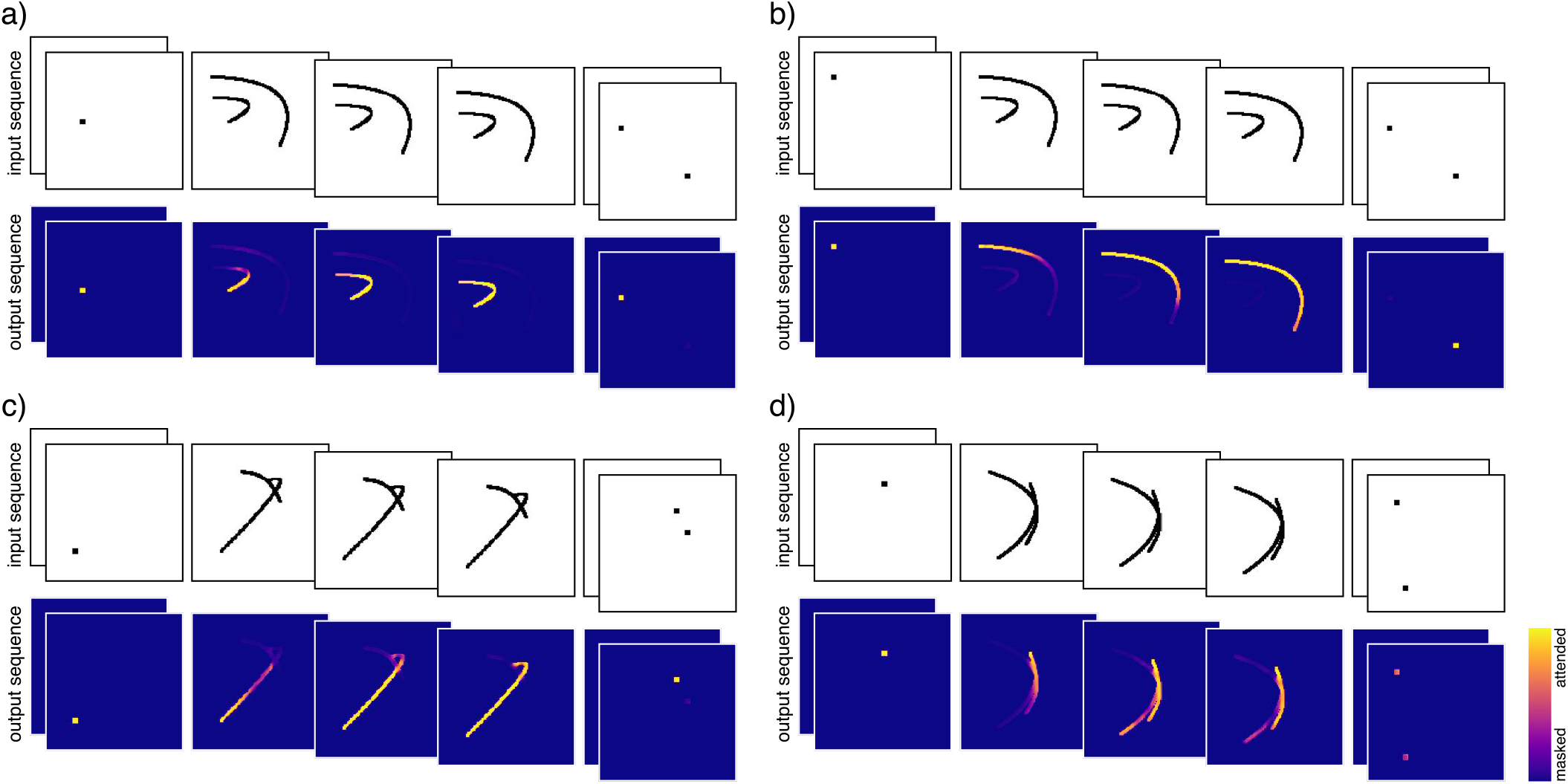
Multiple samples from the curve-tracing experiment. **a-b)** Stimuli and output attention map for a short and a long curve seems to indicate presence of a attentional spread. **a)** Attention propagation on the short curve versus **b)** attention propagation on the long curve. **c** Example of an ambiguous sample where the model makes the correct attention and decision. **d** Example of an ambiguous sample where the model fails to trace the correct curve and make the final decision.

### Single-task training

Our model performs reasonably well when trained on single tasks, if both target class and target attention maps are provided during training. Surprisingly, for MNIST compositions, it is also able to perform object recognition, spatial cueing, and pop-out tasks, even when trained purely on classification (Fig. 17). Unsurprisingly, in our limited grid search, we did not find an effective hyper-parameter regime for the abstract visual cue experiment that achieves good classification accuracy and retrieves a sensible attention map. Our hypothesis is that it is far too challenging for the network to separate the arrow from the digits as different entities, infer the correct sequence of attention, and perform digit classification. With this in mind, we re-designed the sequenced such that the arrow is shown first, then the combination of the arrow and digits, and finally only the digits are left (Fig. 17 d). This new sequence helps the network to use temporal information to distinguish arrow and digits as different entities, and learn to perform the task correctly (Fig. 17 d). The results shown below are from scaled down networks, with no hyper-parameter tuning, and purely to test whether the architecture can in principle learn to perform single tasks.

**Figure 17.**
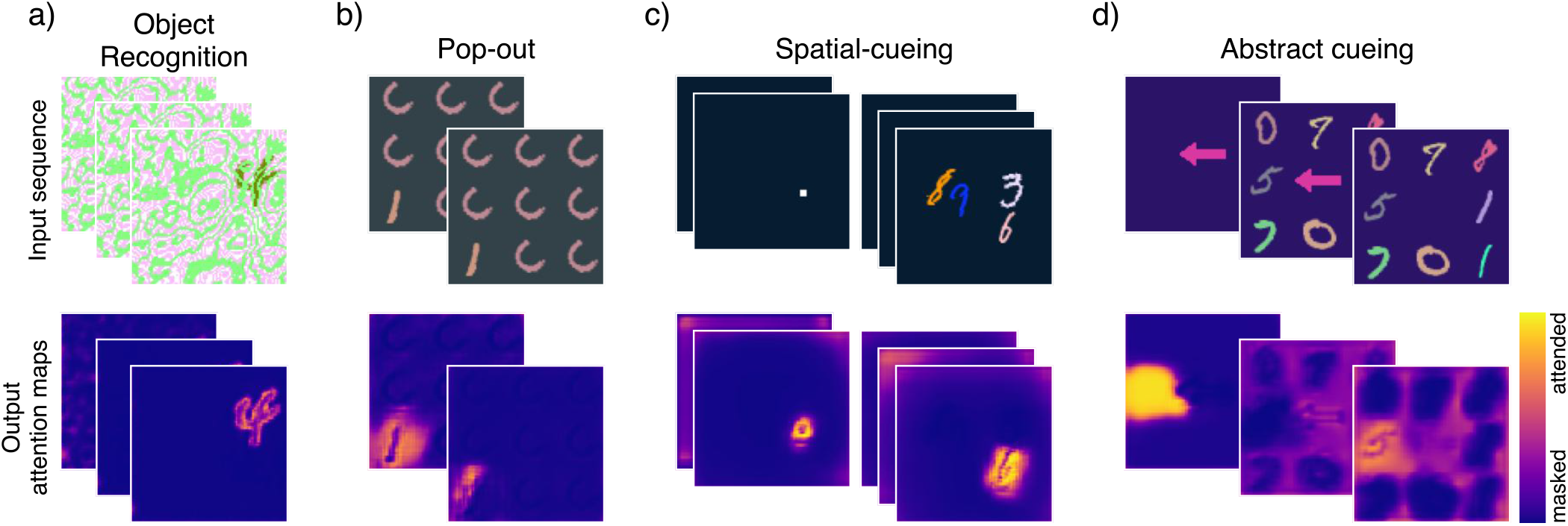
Single-task training on MNIST compositions. Output attention maps for distinct networks separately trained solely on classification for different input compositions. **a)** Classification of digits placed between static background and foreground. **b)** Classification of the “pop-out” digit. **c)** Classification of spatially cued digit. **d)** Classification of the digit which is pointed to by an arrow. Here we use temporal segregation to help the network infer abjectness.

Similarly, we trained a single model on object recognition for the COCO compositions, again purely trained on classification (Fig. 18). As with the original experiment, we used dynamic background for training on the object recognition task (Fig. 18 a) which results in a good output attention map without ever having seen target segmentation maps. We also tested how the network would compare if trained on still images but with similar statistics (Fig. 18). The results indicate that the model trained on dynamic composition does better on classification and is able to output a more recognizable attention map (Fig. 18). The results shown here are from scaled down networks, with no hyper-parameter tuning, and purely to test whether the architecture can in principle learn to perform object recognition and that motion seems to be a helpful signal for learning.

**Figure 18.**
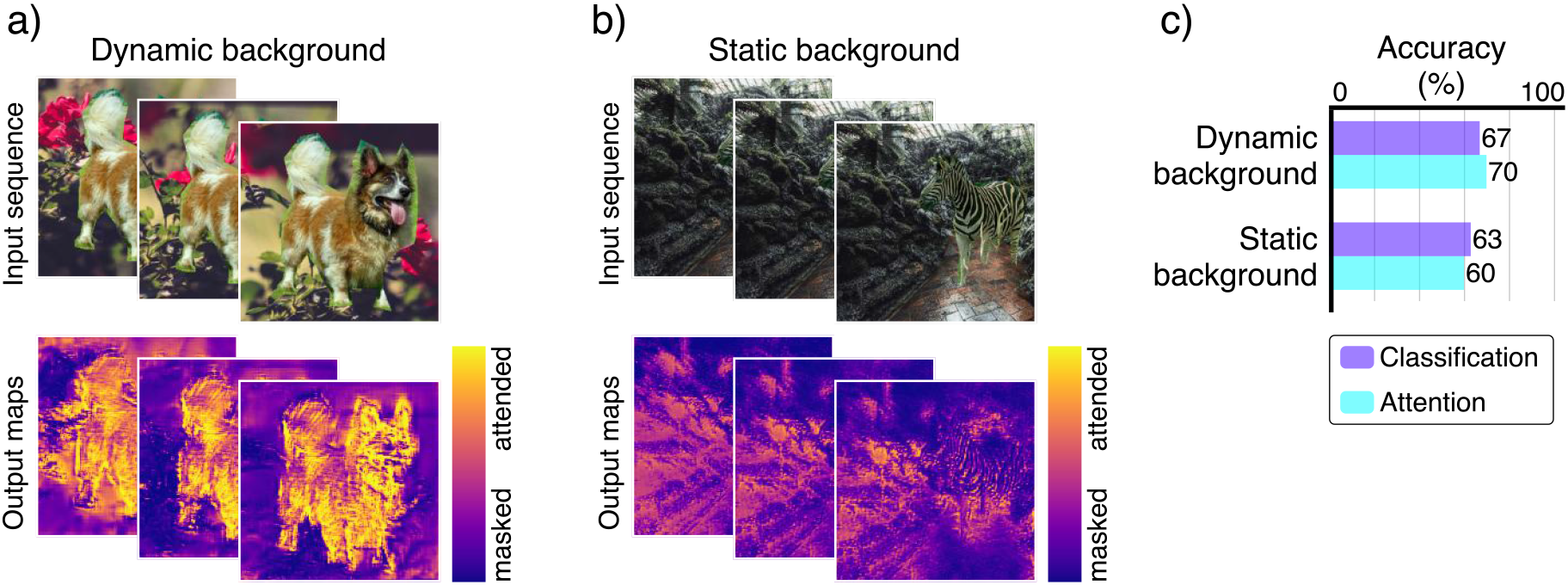
Single-task training on COCO compositions. Output attention maps for networks separately trained solely on classification for different input compositions. Classification of animals placed on **a)** dynamic background and **b)** static background. **c)** our primary results point to the hypothesis that motion is a helpful signal for object recognition and separation.

### Multi-modal top-down visual search

We also trained a separate model based on our architecture on a top-down visual search for class and color using the FashionMNIST dataset [Xiao et al., 2017]. The input prompt, therefore, includes both the target class and color to search for. In this experiment, we also included multiple instances (Fig. 19 left) as well as no match (Fig. 19 right).

**Figure 19.**
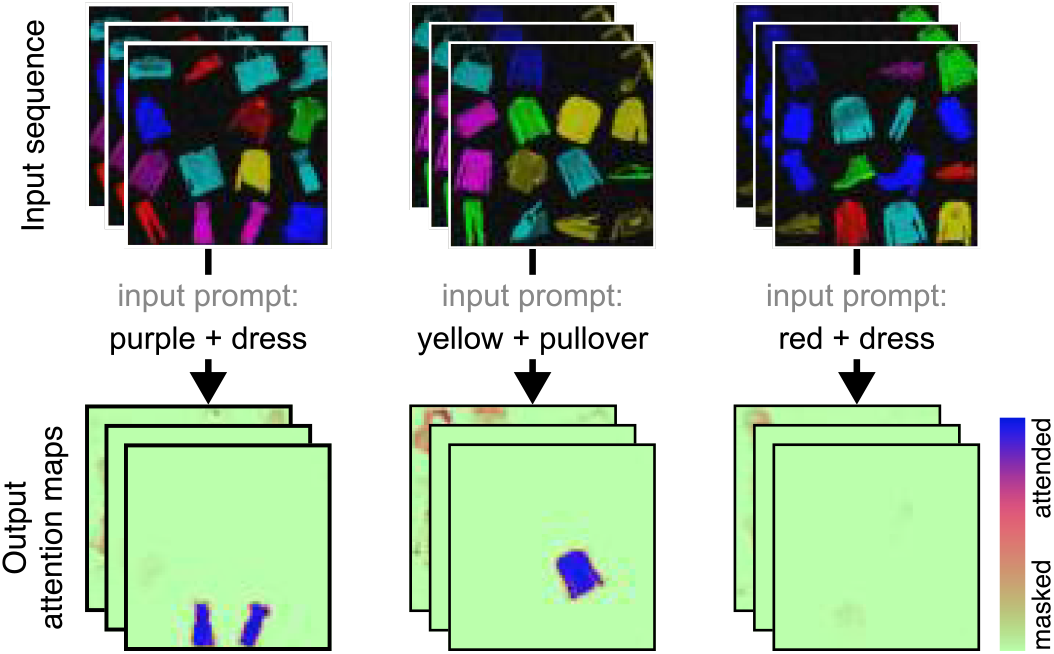
Top-down multi-modal visual search.

## Notes

### Competing Interest Statement

The authors have declared no competing interest.

### Summary of Updates

beautified figures, better writing, and experiments

